# miRNA-mediated expression of RBFOX2 governs the splicing transition from progenitors to neurons in the developing brain

**DOI:** 10.1101/2024.09.20.614071

**Authors:** Stephan Weißbach, Hristo Todorov, Azza Soliman, Laura Schlichtholz, Sophia Mühlbauer, Lea Zografidou, Sarah Lor-Zade, Anna Wierczeiko, Dennis Strand, Susanne Strand, Tanja Vogel, Martin Heine, Susanne Gerber, Jennifer Winter

## Abstract

Alternative splicing is a crucial component of neuronal differentiation, yet the mechanisms that regulate splicing transitions during embryonic brain development remain incompletely understood. Here, we identify a post-transcriptional mechanism that times the expression of the splicing factor *Rbfox2* during neurogenesis. RBFOX2 is normally expressed at low levels in neural progenitor cells (NPCs) and becomes upregulated in newborn neurons where it promotes neuronal differentiation. Unexpectedly, premature expression of *Rbfox2* in NPCs of the embryonic mouse neocortex blocked their differentiation into neurons rather than promoting it. Genome-wide analysis revealed widespread alternative splicing changes enriched for NDD genes and associated with a hybrid NPC- and neuron-like splicing pattern that significantly deviates from the normal splicing developmental trajectory. Remarkably, premature *Rbfox2* expression induced the inclusion of validated target exons that are otherwise repressed by PTBP2 pointing to an antagonistic splicing relationship. Integrative scRNA-seq analysis confirmed a negatively correlated expression between these two RNA-binding proteins (RBP) along differentiation pseudotime. Strikingly, we identified the NPC-specific miRNA 92a-3p as a regulator of the *Rbfox2* expression switch: expression of miR-92a reduced RBFOX2 levels and reversed splicing patterns of target genes in vitro, while silencing miR-92a *in vivo* increased RBFOX2 expression in the embryonic cortex. Together, these findings reveal a previously unrecognized miRNA–RBP regulatory axis that ensures the proper timing of NPC-to-neuron splicing transitions in the developing cortex and provide new insights into splicing dysregulation as a contributing factor to the emergence of neurodevelopmental disorders.

**Graphical Abstract:** 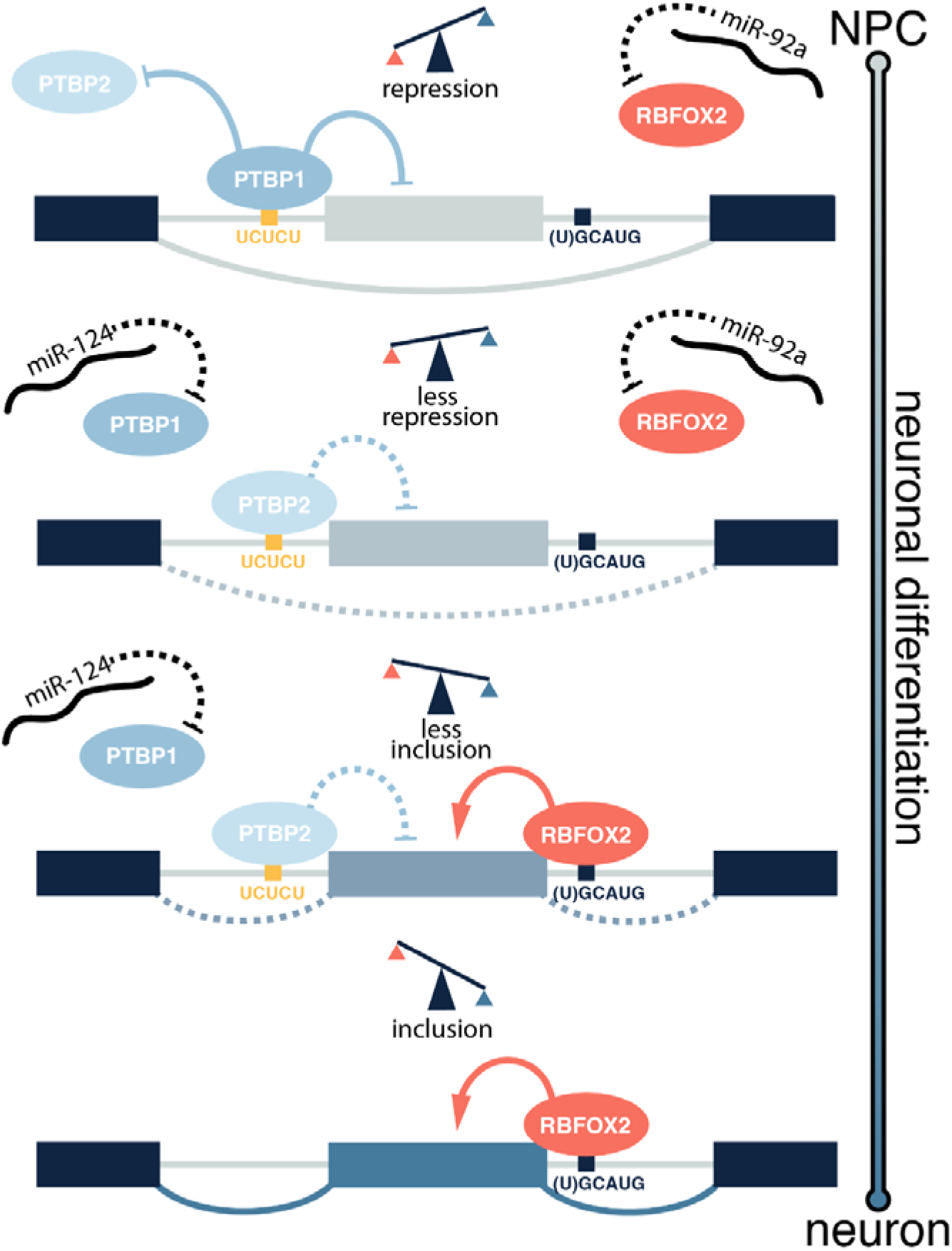

Graphical Abstract Schematic representation of the proposed splicing regulation for the transition of undifferentiated NPCs to neurons in the developing cerebral cortex.

## Introduction

Despite the presence of only approximately 22,000 protein-coding genes^1^, the murine neocortex and hippocampus exhibit a considerable diversity of neurons, amounting to 388 transcriptomic cell types, as reported by the Allen Brain Consortium^2^. A key factor enabling this diversity and complexity is alternative splicing (AS), a tightly regulated process that governs the production of multiple mRNA isoforms from a single gene. Intriguingly, nearly every multi-exon gene in the human genome can undergo alternative splicing, generating numerous protein isoforms with distinct structural and functional properties^3^. This vast molecular diversity expands the proteome’s complexity without necessitating a proportional increase in the genome’s size. Within the nervous system, AS is a pervasive phenomenon, with various neural genes subject to this regulation^4^. Overall, AS has a higher prevalence across different species and is more conserved in the brain compared to other tissues^5,6^.

Specifically, numerous changes in the splicing landscape occur in the developing brain with high temporal precision^4,7^. Among the processes that require correct splicing are neuronal migration, neuronal differentiation^8^, axonogenesis^9^, and synaptogenesis^10^. Notably, the significance of correct splicing is not confined to the developing brain alone. In the adult brain, different neuronal types exhibit distinct splicing patterns, further underscoring the importance of alternative splicing in maintaining the brain’s functional diversity^11,12^.

During embryonic development, neural stem cell differentiation requires precise, extensive AS switches to create neuron-specific splicing patterns. Many of these splicing events alter key protein domains, particularly in proteins of the cytoskeleton^8^. Various RNA-binding proteins (RBPs) regulate the transition from neural stem and progenitor cells to neurons by altering the splicing pattern of target genes, therefore their temporal expression needs to be strictly regulated. Here, microRNAs (miRNAs) that suppress gene expression post-transcriptionally by binding to the 3’ untranslated region of target mRNAs^13–16^ represent one important mechanism for mediating stage and cell-type-specific levels of RBPs. For instance, PTBP1 is expressed only in neural stem and progenitor cells, however, it is strongly repressed by miR-124-3p in neurons^17^. In contrast, other prominent RBPs, including NOVA, PTBP2, MBNL2, and RBFOX, are predominant in neurons. These RBPs act in both synergistic (e.g., NOVA and RBFOX) and antagonistic relationships (e.g., PTBP1 and RBFOX)^18,4,8^.

The RBFOX-family consists of three paralogs, namely, RBFOX1 (FOX-1 or A2B1), RBFOX2 (FOX-2 or RBM9) and RBFOX3 (FOX-3, HRNBP3, or NeuN). The association of mutations in human *RBFOX1* and *RBFOX2* with intellectual disability, autism, and other neurodevelopmental disorders, as well as the specific neurological phenotypes observed in *Rbfox* knockout mice, highlights the important functions of these proteins during neurodevelopment^19–26^. All RBFOX-proteins have a single, highly conserved RNA recognition motif (RRM) that binds to the target sequence (U)GCAUG harbored in the premature mRNA and initiates position-dependent splicing. RBFOX binding in the intronic region upstream of an alternatively spliced exon promotes its exclusion from the mature mRNA, whereas downstream binding leads to inclusion^27^. Moreover, in humans, all RBFOX-proteins share the position-dependent splicing mechanism and the RRM, which is identical in RBFOX1 and RBFOX2 and almost identical in RBFOX3. Therefore, the RBFOX family can act in a functionally redundant manner^28^. However, the temporal expression patterns of these proteins diverge significantly, exhibiting variations across different tissues and cell types^2,11^. In the developing brain, RBFOX proteins are either lowly expressed or not present in neural stem cells, but they are highly expressed in neurons, which is in agreement with their role in promoting neuronal differentiation. Studies aiming at identifying splicing changes during neural stem cell differentiation have compared alternative exon usage between NPCs and neurons by RNA-sequencing^8,29^. While these studies have identified numerous splicing changes that are presumably driven by RBFOX binding, the exact role of RBFOX during neuronal differentiation in the embryonic brain has not been elucidated yet.

In the current study, we identified extensive AS changes upon premature *Rbfox2*expression in the embryonic neocortex, specifically related to genes that are important for neuronal differentiation and localization. Remarkably, we observed a mixed neuronal and NPC-like splicing pattern that significantly deviated from the normal cortical splicing trajectory. In line with this, *Rbfox2* expression increased the percentage of neural stem and progenitor cells and blocked differentiation into neurons on the cellular level. Since a precise and tight control of RBFOX2 is essential for proper neuronal differentiation in the embryonic neocortex, we investigated whether miRNAs might repress *Rbfox2* expression in NPCs. Indeed, we observed that silencing the NPC-specific miR-92a-3p in the embryonic neocortex was associated with a concomitant increase in *Rbfox2* expression. Conversely, miR-92a-3p overexpression resulted in reduced RBFOX2 transcript and protein levels, as well as a reversal of the splicing outcome in RBFOX2-targeted genes.

## Results

### RBFOX2 is predominantly expressed in neurons

The RBFOX protein family exhibits distinct temporal expression patterns, also varying by cell type. For instance, RBFOX1 and RBFOX3 are expressed in mature neurons and exclusively in the cortical plate in the embryonic cerebral cortex^30^. RBFOX2 shows early expression in Purkinje cells in the cerebellum^31^. To elucidate the composition of RBFOX proteins in the mouse embryonic cerebral cortex, we co-stained sections for RBFOX1 and RBFOX2. This confirmed RBFOX2’s presence was not limited to the neuron-specific cortical plate, but it was also expressed in the intermediate and ventricular (VZ)/subventricular (SVZ) zones (Figure 1a). In contrast, RBFOX1 expression was restricted to the cortical plate/subplate. While this broader expression pattern of RBFOX2 compared to RBFOX1 was evident, not all cells in the cerebral cortex showed positive RBFOX2 staining. Especially in the VZ/SVZ, where radial glia cells and basal progenitors reside at this developmental stage, the majority of cells were RBFOX2 negative (Figure 1a). This prompted us to investigate the temporal expression of RBFOX2 using an *in vitro* neuronal differentiation model. Here, we observed low *Rbfox2* mRNA expression in radial glia-like cells and early neural progenitor cells (NPCs). However, *Rbfox2* expression increased significantly during neuronal differentiation, peaking in late NPCs and young neurons (Figure 1b). Likewise, *Rbfox2* was only weakly expressed in NPCs derived from the embryonic cerebral cortex, while it was strongly upregulated in cortical neurons at both RNA and protein levels (Figure 1c, d).

**Figure 1.**
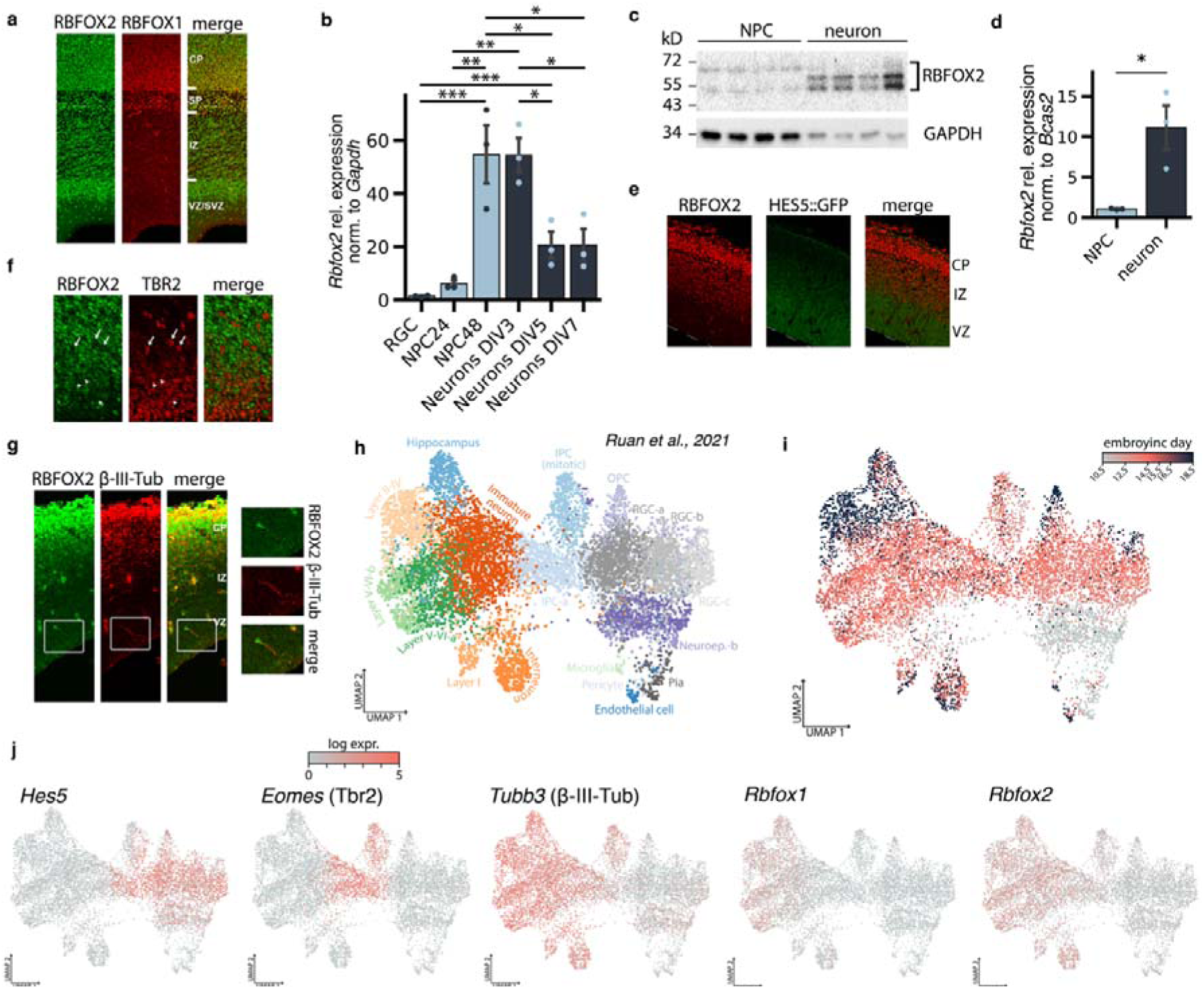
Rbfox2 expression was upregulated in newborn neurons. **a)** Cryosections of E17.5 cerebral cortex were co-stained with antibodies against RBFOX1 and RBFOX2. Expression of RBFOX1 was restricted to the cortical plate/subplate whereas RBFOX2 was additionally expressed in cells of the ventricular zone (VZ)/subventricular zone (SVZ) and intermediate zone (IZ). **b)** To determine the expression of Rbfox2 during neuronal differentiation, embryonic stem cells were differentiated into radial glia cells (RGCs), NPCs, and neurons and subjected to RT-qPCR with Rbfox2-specific primers. Rbfox2 expression increased strongly in late NPCs and early neurons and decreased again in late neurons. **c)** Western blot analysis with RBFOX2-specific antibodies illustrates the increase in RBFOX2 expression in neurons compared to NPCs. Note that due to alternative splicing, several RBFOX2 isoforms exist which differ between NPCs and neurons. **d)** RT-qPCR showed increased Rbfox2 expression in neurons compared to NPCs. **e)** Cryosections of the E14.5 cerebral cortex derived from HES5::GFP transgenic mice were immunostained with RBFOX2 antibodies. RBFOX2 was predominantly expressed in differentiating GFP^-^cells. **f)** and **g)** Cryosections of the E17.5 cerebral cortex were co-immunostained for RBFOX2 and the basal progenitor marker TBR2 (f) or RBFOX2 and the newborn-neuron marker β-III-Tubulin (g). RBFOX2 was expressed in β-III-Tubulin^+^ neurons and absent from TBR2^+^ basal progenitors. **h)** UMAP representation of publicly available scRNA-seq data of the developing murine neocortex (GSE161690)^32^, colors represent cell-type assignment. IPC: intermediate progenitors, Neuroep.: Neuroepithelial cells, OPC: oligodendrocyte progenitor cells i) The scRNA data includes cells from E10.5, E12.5, E14.5, E15.5, E16.5, and E18.5. j) Expression of marker genes used in the immunostainings (e,f,g), and Rbfox1/2. *p<0.05, **p<0.01, ***p<0.001, ANOVA followed by Tukey’s post-hoc test in b, t-test in d.

To further corroborate these data, we analyzed RBFOX2 expression in the embryonic cerebral cortex *in vivo* by immunostaining. To this end, we made use of a transgenic mouse model (HES5::GFP) expressing GFP exclusively in Notch-signaling neural progenitor cells. In agreement with the *in vitro* data, RBFOX2 expression was restricted to HES5 negative cells, that were presumably in a more differentiated state compared to HES5-expressing NPCs (Figure 1e). Co-stainings with the basal progenitor marker TBR2 (EOMES) confirmed the absence of RBFOX2 from neural progenitor cells. Only cells with low TBR2 signal exhibited RBFOX2 expression, suggesting that these cells were currently differentiating into neurons (Figure 1f). In contrast, RBFOX2 was highly expressed in newborn neurons in all zones as revealed by co-staining with the neuron marker β-III-Tubulin (TUBB3) (Figure 1g).

To validate these findings with a single-cell resolution, we integrated a publicly available data set of the developing mouse neocortex at six developmental time points from E10.5 through E18.5^32^. These data robustly captured cell type heterogeneity and developmental trajectories of the mouse cerebral cortex with an enrichment of neuroepithelial cells (NECs) at E10.5, transitioning through radial glia cells (RGCs) and intermediate progenitors (IPCs) peaking at E12.5-E14.5 and finally differentiating to immature and cortical layer neurons (Figure 1h,i, Supplementary Figure S1a). In agreement with our results, *Rbfox1* was restricted to mature neurons, however, *Rbfox2* exhibited a broader expression pattern (Figure 1j). Specifically, no expression was detected in NECs and *Hes5*-positive RGCs, and only low levels were present in *Eomes (Tbr2*)-positive IPCs. In contrast, *Rbox2* became up-regulated in *Tubb3*-expressing immature neurons and retained high levels in cortical neurons (Figure 1j, thus confirming results from our immunostaining experiments. In summary, RBFOX2 was weakly expressed in neural stem and progenitor cells and highly upregulated during neuronal differentiation, highlighting its potential role in mediating the neuron-specific splicing switch.

### Premature *Rbfox*2 overexpression disrupts the splicing landscape of the developing cortex

To study RBFOX2’s impact on splicing during embryonic corticogenesis, we used *in-utero* electroporation to overexpress *Rbfox2* in the mouse cerebral cortex at E13.5 and performed RNA-sequencing at E15.5 (Figure 2a). At E13.5, the developing cortex is dominated by progenitor cells (Supplementary Figure S1a), which do not express *Rbfox2* normally (Figure 1j). This resulted in a significant increase in *Rbfox2* gene expression (p_adj_ < 0.0001, log2fold = 5.754, Figure 2b), 268 differentially expressed genes (Supplementary Figure S1b,c, Supplementary Data 1), and substantial alternative splicing changes compared to controls (Figure 2c). However, most alternatively spliced genes did not show altered expression levels. Only 30 genes were alternatively spliced and differentially expressed at the same time (Supplementary Figure S5a), and no significant expression changes related to nonsense-mediated decay were observed despite potential frameshifts (Supplementary Figure S5b).

**Figure 2.**
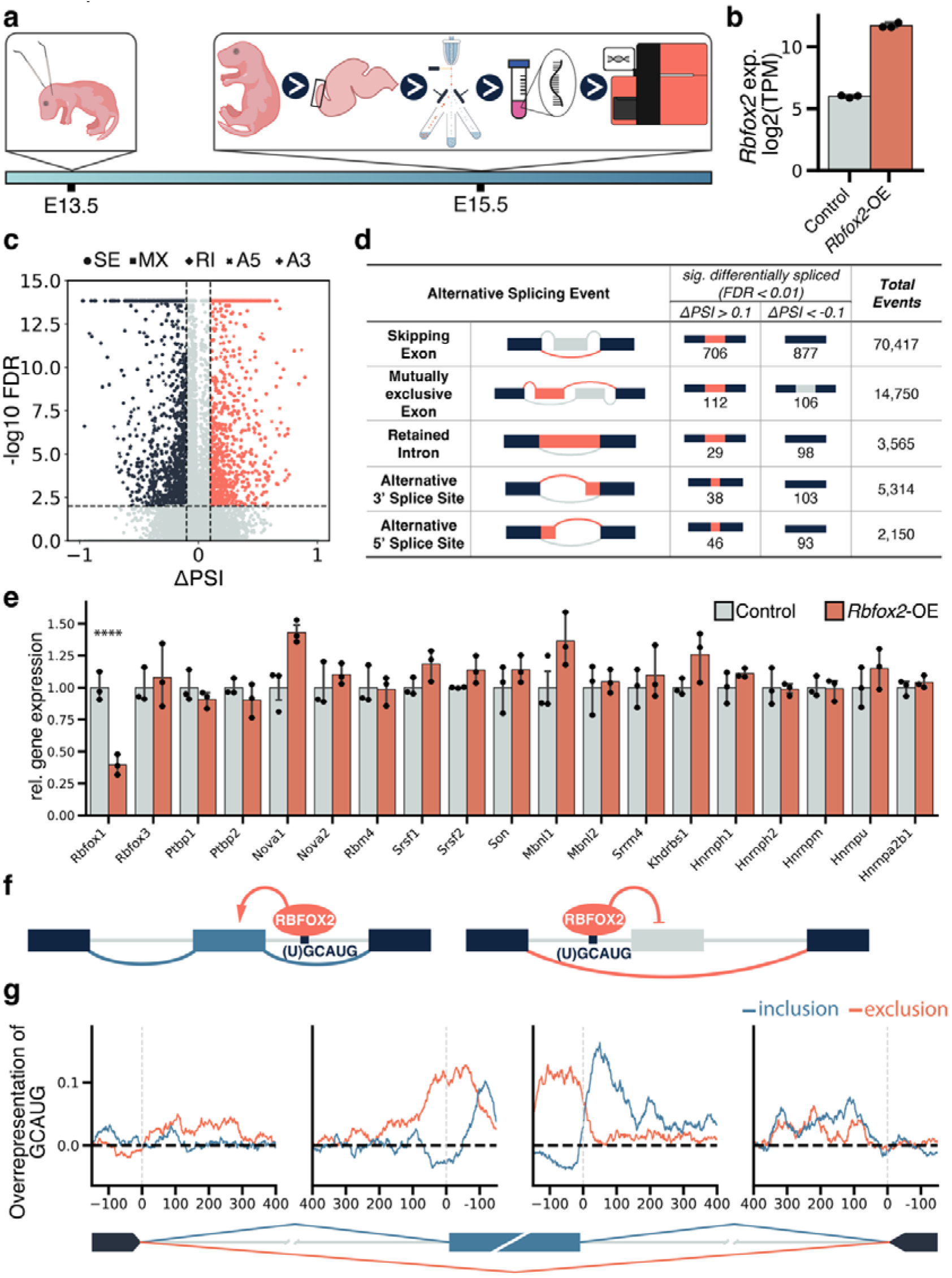
Rbfox2 overexpression changes the splicing pattern of the developing cortex. **a)** Experimental timeline, Rbfox2 overexpression was induced in embryonic mouse neocortex at E13.5 by in-utero electroporation, and bulk RNA-sequencing was performed at E15.5. **b)** The electroporation of Rbfox2 constructs increased the mRNA abundance of Rbfox2 in the cortex significantly. **c)** The volcano plot visualizes alternative splicing events. The x-axis shows delta percentage spliced in (ΔPSI), and the y-axis shows -log10 false discovery rate (FDR) values. Significantly spliced-in events are in orange, spliced-out events are in blue, and non-significant events are in grey. Shapes denote event types: circles for skipping exons, squares for mutually exclusive exons (MX), diamonds for retained introns (RI), and crosses for alternative A5 and A3 splice sites. **d)** The table provides an overview of the 2208 significant alternative splicing events. Significance was defined as FDR < 0.01 and either ΔPSI > 0.1 or ΔPSI < -0.1. **e)** Normalized gene expression levels of key splicing factors important in the CNS, upon Rbfox2 overexpression compared to control. Values for each gene are normalized to the mean of the control group. Data are shown as mean ± standard error of the mean *** p<0.001, ****p<0.0001, Wald test from DESeq2 in g. **f)** Schematic representation of the position-dependent splicing of skipping exons by RBFOX2. **g)** Relative abundance of the RBFOX2-motif GCAUG in the vicinity of alternatively spliced exons and their flanking exons.

We detected 2,208 significant alternative splicing events (FDR < 0.01, |ΔPSI| > 0.1), predominantly corresponding to skipped exons (∼71%, Figure 2d). These changes can be attributed directly to RBFOX2 as the expression of other known neurodevelopmental splice factors was unchanged (Figure 2e). The only exception was a significant reduction of *Rbfox1* (p_adj_ < 0.0001, log2fold = -1.342). Furthermore, we observed a significant overlap between skipping exon events in our data with *Rbfox* triple knock-out (t*Rbfox*-KO) samples^33^ (fold enrichment ≈ 9.833, p-value < 0.0001; Supplementary Figure S3b, Supplementary Data 2). The inclusion levels of cassette exons present in both data sets were strongly negatively correlated (R = - 0.6624, p < 0.0001), indicating that *Rbfox2-*OE induced the opposite splicing regulation compared to t*Rbfox*-KO, thus strengthening the biological validity of our results (Supplementary Figure S3a).

Remarkably, alternatively spliced genes in our analysis were significantly enriched for high-confidence neurodevelopmental disorder (NDD) genes^34^ (224 out of 1317 genes, fold enrichment ≈ 1.358, p < 2e-07, Supplementary Data 3, Supplementary Figure S4).

RBFOX proteins preferably bind to the (U)GCAUG^35,36^ motif with position-dependent effect: upstream intronic binding represses exon inclusion while downstream binding enhances exon inclusion (Figure 2f)^37^. Therefore, Wwe screened pentameric motifs in the 150 nt intronic region flanking alternatively spliced cassette exons. As expected, the canonical RBFOX motif GCAUG was enriched downstream of included exons (z-score: 9.4) and upstream of repressed exons (z-score: 6.8), with related motifs UGCAU and CAUGC showing similar patterns (Supplementary Figure S5c). Positional analysis of the canonical GCAUG motif revealed that exon skipping was promoted by motifs located ∼100 nt upstream and on the exon itself, while exon inclusion was associated with GCAUG occurrence ∼150 nt downstream, consistent with previous reports on RBFOX2’s position-dependent splicing activity (Figure 2g)^38^. Moreover, we detected highly enriched poly CU-motifs upstream of included exons, associated with PTB proteins^39^ that initiate exon exclusion upon binding^40^. Analysis of publicly available RBFOX2-RNA interaction data (iClip)^38^ confirmed that most enriched motifs around alternatively spliced exons are direct RBFOX2 binding sites (Supplementary Figure S5d, e).

To gain a perspective on the global splicing pattern induced upon *Rbfox2*-OE, we next integrated the previously published RNA-sequencing data from Weyn-Vanhentenryck et al. obtained at different developmental stages of the mouse cerebral cortex (E14.5 to P30)^41^, as well as RNA-sequencing from FACS-sorted NPCs and neurons^29^. The hierarchical clustering of splicing events that were significant both in our data and in the previous studies revealed that our control group clustered with the FACS-sorted neurons as well as with the E14.5 and E16.5 cortical samples. In contrast, the *Rbfox2*-OE group did not closely cluster with any of the embryonic samples or NPCs/neurons, pointing to an abnormal splicing pattern (Supplementary Figure S2).

To further investigate the impact of *Rbfox2*-OE on the splicing landscape during neurogenesis, we applied a principal component analysis (PCA). We observed a bell-shaped trajectory indicative of the expected splicing changes throughout cortical development (Figure 3a)^42^. *Rbfox2*-OE resulted in a substantial deviation from this expected developmental trajectory. Specifically, on the first principal component (PC1), *Rbfox2*-OE exhibited a shift towards the splicing pattern associated with mature neurons, while the shift on PC2 was towards undifferentiated NPCs. To identify the main splicing events that explained the observed PCA pattern, we calculated the loadings of the original variables on PC1 and PC2 (Figure 3b). Interestingly, genes with the highest loadings are associated with neurodevelopmental diseases. For instance, *Shank1* and *Dlgap4* are related to autism spectrum disorder^43,44^, whereas *Dlg2* has been linked with schizophrenia^45^.

**Figure 3.**
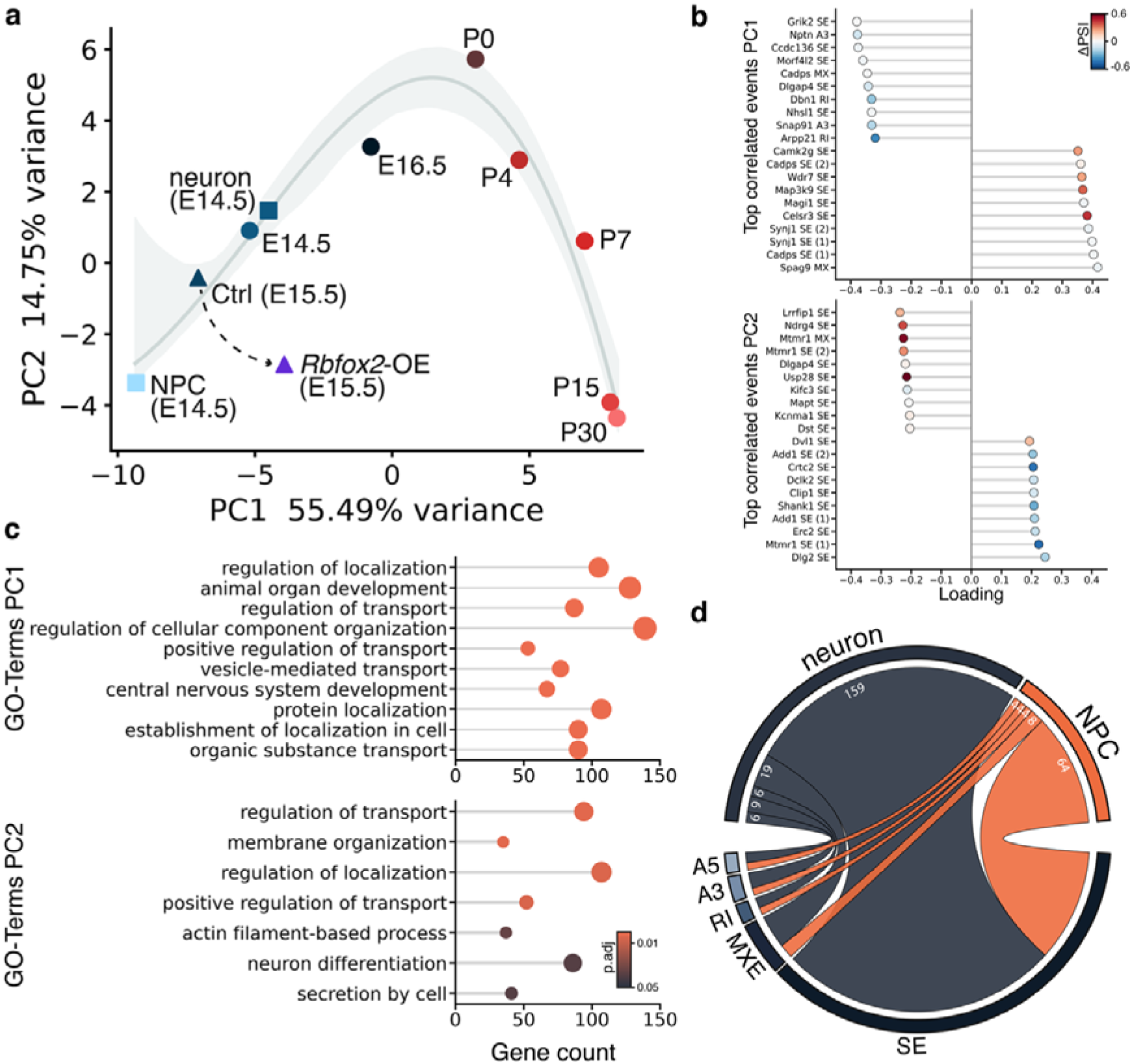
Developmental regulation of alternative splicing by RBFOX2. **a)** Principal Component Analysis (PCA) of splicing patterns using SplicePCA^42^ across embryonic and postnatal brain development. Blue color dots represent embryonic and red dots denote postnatal stages. Shapes indicate the source of the data used. Circles correspond to E14.5 to P30 samples from Weyn-Vanhentenryck et al.^41^, and squares represent NPC and neuron samples from Liu et al. ^47^. The developmental trajectory (gray line with 95% confidence intervals) was fitted using spline regression with bootstrap resampling. Rbfox2 overexpression samples (triangles) deviate from the normal developmental trajectory. **b)** Loading analysis for principal component (PC) 1 and PC2, highlighting the top 10 positively and top 10 negatively correlated splicing events for each PC. Color gradient represents the ΔPSI values for Rbfox2-OE compared to control samples. **c)** Gene Set Enrichment Analysis (GSEA) performed on splicing events for PC1 and PC2 using loadings on each component as effect sizes. **d)** Chord plot showing the association between significant splicing events in Rbfox2-OE vs. control samples compared with splicing events in neurons vs. NPCs obtained using the data from Liu et al. ^47^.

Furthermore, a Gene set enrichment analysis (GSEA) of gene ontology (GO) terms (Figure 3c, Supplementary Data 4) revealed that PC1 was associated with higher-level GO terms such as “regulation of transportation”, “animal organ development”, and “central nervous system development”, reflecting the separation of samples across a broad developmental window from E14.5 to P30.

Moreover, neuron differentiation was among the significant terms associated with PC2, where the *Rbfox2*-OE samples were more similar to an NPC splicing pattern. When analyzing the splicing events that significantly contributed to this GO term, we identified several developmentally relevant genes. For instance, *Clasp2*, a protein involved in microtubule dynamics, is known to play a role in neuronal migration and axonal growth during cortical development^46^. We detected an alternatively spliced exon in *Clasp2* (chr9:113,691,503-113,691,530, Supplementary Figure S6b) that showed a significantly higher inclusion in the *Rbfox2* overexpression group (FDR < 0.0001, ΔPSI = 0.118). Interestingly, we observed a similar increase in the same exon when analyzing the splicing changes between NPCs and neurons (FDR < 0.01, ΔPSI = 0.118).

The observed mixed splicing pattern upon *Rbfox2*-OE was not only characterized by the deviation from the expected developmental trajectory (Figure 3a) but also by the type of splicing events. When comparing the significant splicing events upon *Rbfox2*- OE with those of neurons vs. NPCs from the Liu et al. data^47^, we found an overlap of 283 significant events. Strikingly, 70% of the events corresponded to a neuron-like splicing and 30% of events represent NPC-like splicing patterns (Figure 3d). Several of the overlapping genes have been reported previously to undergo alternative splicing during important developmental processes associated with NPC proliferation, radial migration of neurons, and neuronal differentiation^8^. Genes with a neuron-like splicing included *Clasp2*, *Dctn1*, *Kif2a*, *Mast2*, and *Tpm1*. In contrast, *Add1*, *Clip1*, *Clasp1*, *Dync1i2*, *Gphn*, and *Macf1* were spliced in an NPC-like way (Supplementary Data 5).

### RBFOX2 misexpression impairs neuronal differentiation in the embryonic neocortex

After ascertaining the pronounced effect of *Rbfox2* overexpression on the splicing landscape of the developing neocortex, we next investigated whether this translates into changes in neuronal differentiation and/or migration at the cellular level. Therefore, we electroporated *Rbfox2* constructs into the E13.5 neocortex and analyzed brain sections two days later by immunostaining experiments. We detected significantly more RBFOX2-OE cells in the VZ and llZ. Conversely, we observed significantly fewer RBFOX2-overexpressing cells in the uIZ and the CP (Figure 4a). In line with the observed phenotype of cells detaching from the ventricular surface and subsequently failing to migrate to the cortical plate and accumulating in the lIZ, *Rbfox2-OE* induced alternative splicing of two key constituents of the Reelin pathway, which plays an important role during cortical development by guiding neurons to the correct position in the CP^48^. Specifically, an exon of *Lrp8* encoding the murine-specific eighth LDL receptor type A (chr4:107,705,477-107,705,600, Supplementary Figure S6f) was abnormally included (FDR < 0.0001, ΔPSI = 0.275). Expression of this exon was previously linked to neuronal migration^49^. Additionally, the exon (chr19:27,217,244-27,217,370, Supplementary Figure S6g) of *Vldlr* encoding the calcium-binding EGF domain was repressed following *Rbfox2-*OE (FDR < 0.0001, ΔPSI = -0.201). Repression of the EGF domain alters the binding affinity to ligands, modulating the sensitivity of the receptor and its interacting ligands^50^. Moreover, RBFOX2 could synergistically act with NOVA2, which is known to induce alternative splicing changes in *Dab1*. DAB1 constitutes a switch within the Reelin pathway that is required for neuronal migration^51,52^.

**Figure 4.**
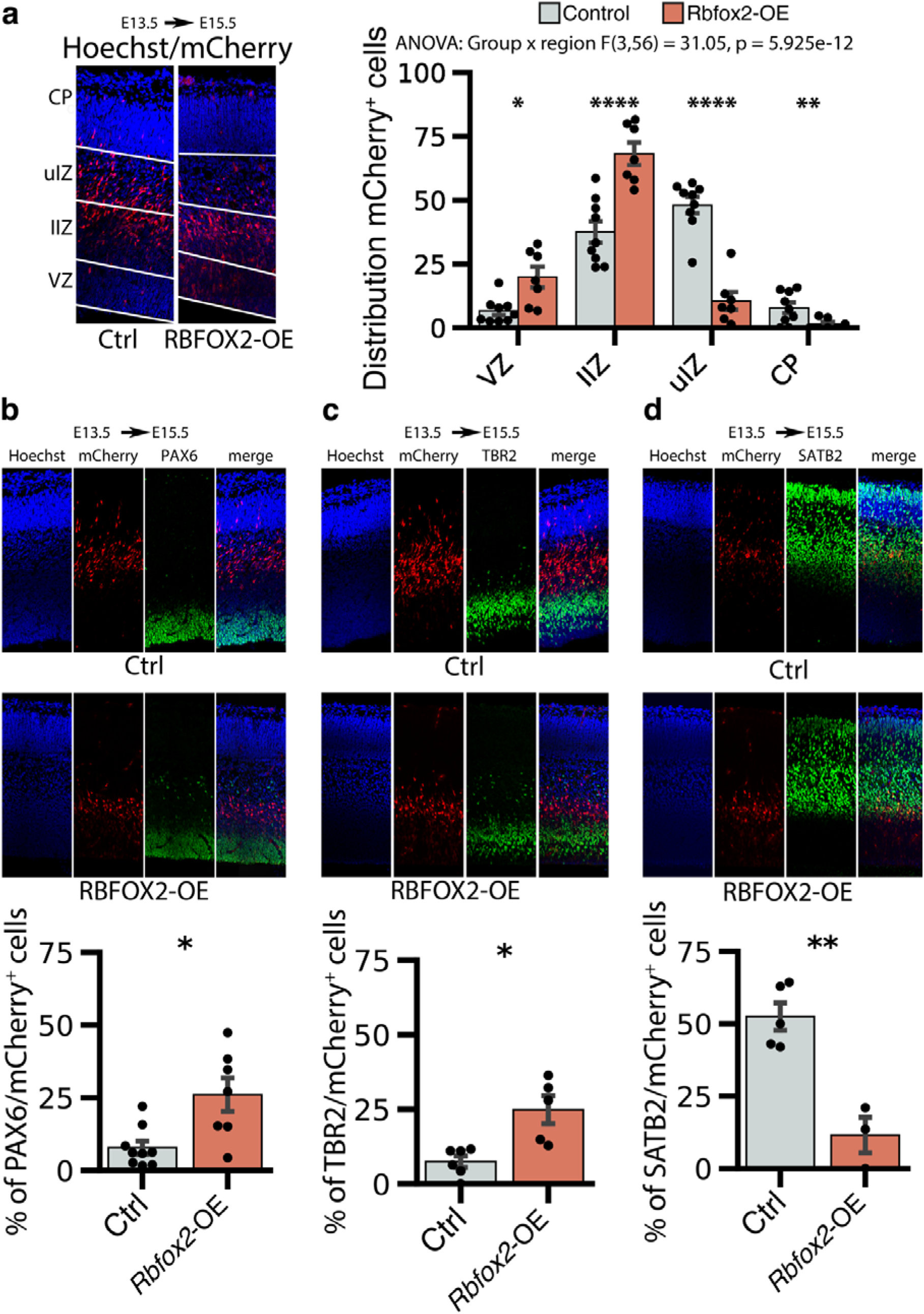
Rbfox2 mis-expression disrupts neuronal differentiation in the neocortex. Rbfox2 expression constructs were co-electroporated with pCAG-mCherry at E13.5. The electroporated brains were analyzed 48 hours later. **a)** Distribution of the electroporated mCherry^+^ cells in the embryonic neocortex. * p<0.05, ** p<0.01, **** p<0.0001, two-way ANOVA followed by pairwise comparison of model means. **b-d)** Cryosections of the electroporated brains were immunostained for markers (green) for radial glia cells (b; PAX6), basal progenitors (c; TBR2) and neurons (d; SATB2). Bar plots show mean values ± standard error of the mean. * p<0.05, ** p<0.01, unpaired t-test in b-d.

To investigate the impact of *Rbfox2* misexpression on neuronal differentiation, we performed immunostainings with antibodies against the marker genes PAX6 (radial glia cells), TBR2 (basal progenitors), and SATB2 (neurons). This revealed a significant differentiation defect in the *Rbfox2* overexpressing neocortex. In particular, the percentage of PAX6^+^ radial glial cells and TBR2^+^ basal progenitors was significantly increased compared to the control group (Figure 4b,c), whereas the percentage of SATB2^+^ neurons was reduced (Figure 4d). This indicates that a correct and fine-tuned expression of RBFOX2 is essential for proper neuronal differentiation in the embryonic neocortex.

### RBFOX2 and PTBP antagonistically regulate shared targets in the splicing transition from NPCs to neurons

The occurrence of the PTBP motifs upstream of significantly spliced-in exons upon *Rbfox2*-OE we previously detected hints at antagonistic splicing activities of RBFOX2 and PTBP (Figure 5a, Supplementary Figure S5c). PTBP proteins bind to pyrimidine-rich sequences located up to 100 nt upstream of alternatively spliced exons^9,53^. Accordingly, we observed a prominent UCUCU peak in the ∼100 nt intronic sequences upstream of exon inclusion events (Figure 5b). Notably, in experiments involving human brain tissue, cell lines, and the mouse cerebral cortex, the upstream occurrence of the PTBP motif was robustly correlated with exon inclusion events. In contrast, in other tissue types, the upstream enrichment of the PTBP motif was associated with exon-skipping^8,53^.

**Figure 5.**
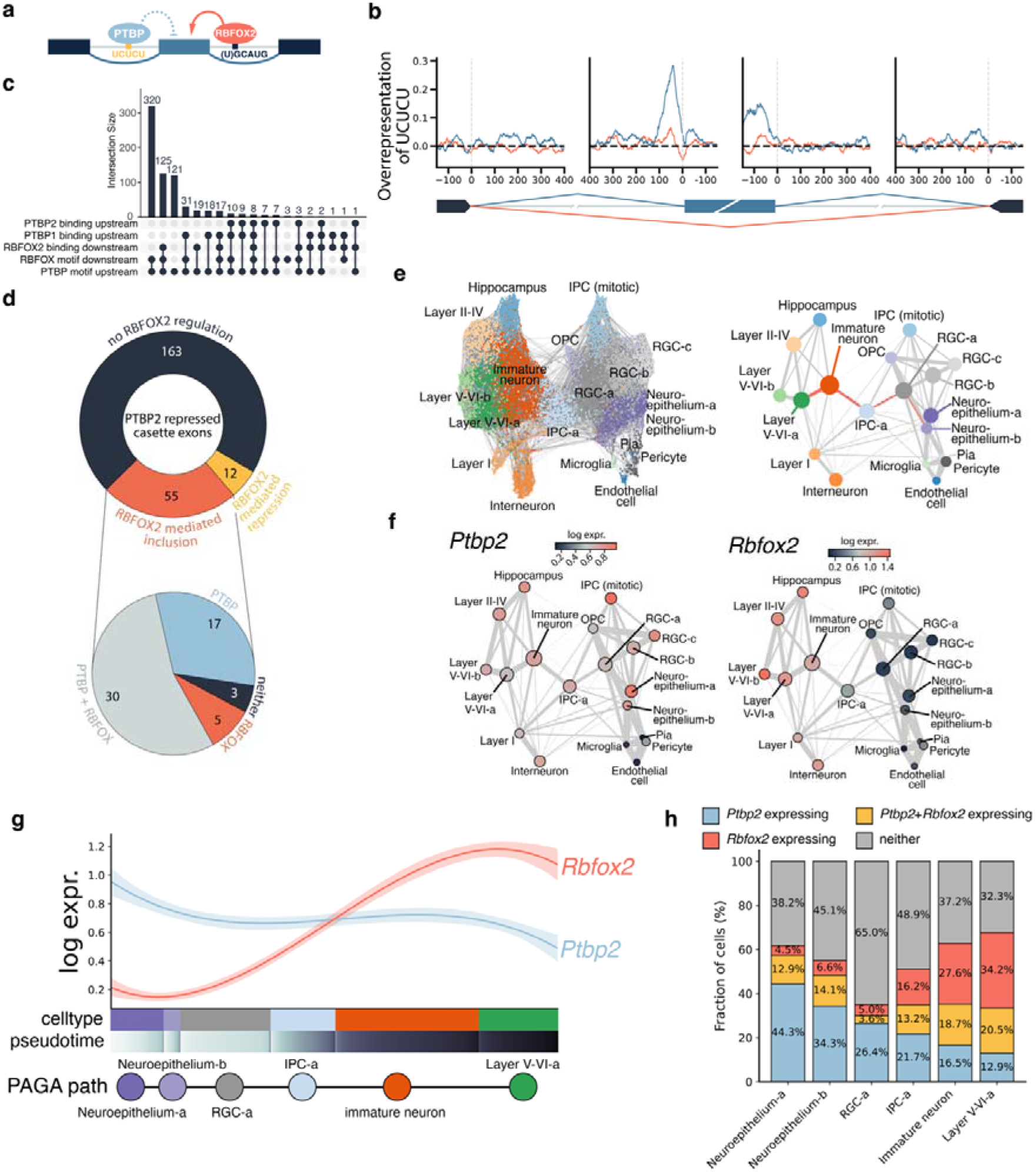
RBFOX2 and PTBP2 regulate common targets in the embryonic neocortex. **a)** Schematic representation of antagonistic splicing regulation between RBFOX2 and PTBP. **b)** Relative abundance of the PTBP motif UCUCU in the vicinity of alternatively spliced exons in Rbfox2-OE vs. control and their flanking exons. Blue lines indicate exon inclusion and red lines correspond to exon skipping events. **c)** UpSet plot showing the overlap of the presence of RBFOX or PTBP motifs in the flanking intronic sequences of alternatively spliced cassette exons (400 nt) as well as direct RBFOX2 and PTBP2 binding interactions as indicated by the presence of peaks obtained from iClip data^38,58^. **d)** Analysis of developmentally regulated cassette exons suppressed by PTBP2 and the overlap with significantly alternatively spliced exons upon Rbfox2-OE. The donut-shaped circle represents 230 neurodevelopmentally relevant PTBP2-repressed cassette exon events. Colors indicate regulation of these exons upon Rbfox2-OE, blue: unregulated, orange: inclusion, yellow: repression. The lower circle summarizes the presence of conserved binding motifs for RBFOX2 and/or PTBP2 for the exons that are regulated in opposite direction by PTBP2 (repression) and RBFOX2 (inclusion). **e)** Force-directed graph embedding and PAGA plot of scRNA-seq data from the developing neocortex^32^, the trajectory of neuronal differentiation is highlighted in orange on the PAGA plot. **f)** Ptbp2 and Rbfox2 expression in different cell types. **g)** Fitted log expression of Rbfox2 (orange) and Ptbp2 (blue) across the neuronal differentiation trajectory (as depicted by the orange path in e, right). Cells are arranged by cell type and differentiation pseudotime. Trend lines were obtained with spline regression, and areas around the fitted lines indicate 95% bootstrap confidence bands. **h)** Fraction of cells expressing either only Ptbp2 (blue), Ptbp2 and Rbfox2 (yellow), only Rbfox2 (orange), or neither of the genes (gray) across the neuronal differentiation trajectory.

These results raise the question of why upstream PTBP binding leads to exon inclusion rather than the expected exclusion in the nervous system. We hypothesized that RBFOX2 might antagonistically overrule the splicing outcome of PTBP and therefore favor exon inclusion in the transition from immature to mature neuronal splicing.

While an interaction between RBFOX and PTBP1 specifically has already been described^8,54–57^, the co-regulation of these RBPs may be less relevant as PTBP1 is primarily expressed in non-neuronal cells^17^, contrary to the expression pattern of RBFOX2 (Figure 6a, Figure 1j, Sup. Fig. 7c).

**Figure 6.**
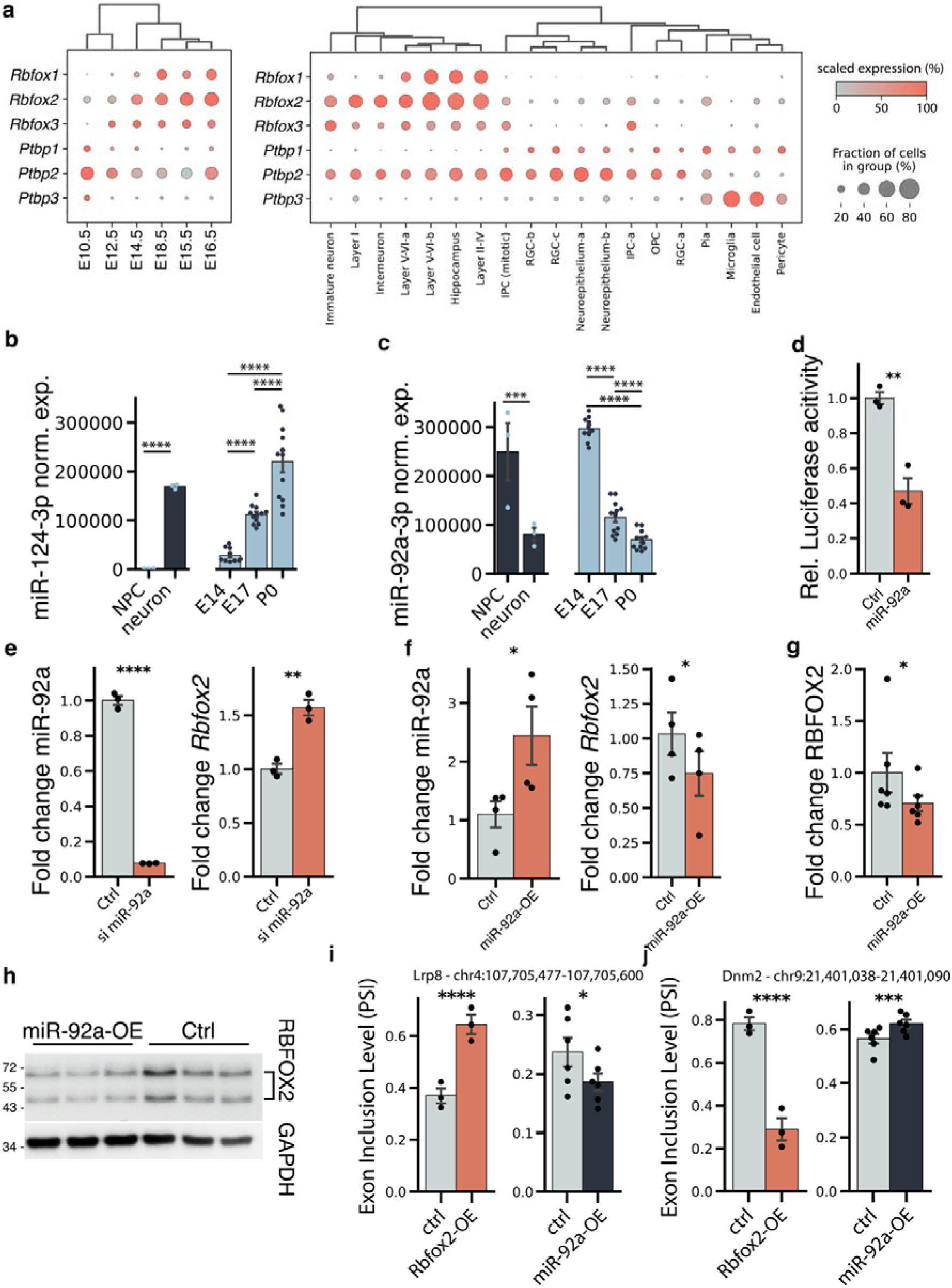
miR-92a-3p regulate RBFOX2 expression and splicing activity. **a)** Dot plot showing the expression of Rbfox1-3 and Ptbp1-3 resolved by developmental stage (left) and cell-type (right) using scRNA-seq from Ruan and colleagues^32^. Dot size indicates the percentage of cells expressing the respective gene, and color indicates scaled expression. **b)** Expression of mir-124-3p known to repress Ptbp1. **c)** Expression of mir-92a-3p predicted to target Rbfox2. Data on miRNA expression shown in c and d were obtained from Todorov et al.^62^. **d)** Luciferase activity in lysates of HEK293 cells transfected with plasmids containing 3’ UTR fragments of Rbfox2 and miR-92a. Fold change of luciferase activity was obtained by calculating the ratio of Renilla luciferase and firefly luciferase activity and then normalizing to the mean of the control. **e)** In vivo knock-down of miR-92a-3p resulted in a more than 10-fold reduction in miR-92a-3p abundance (left side), and subsequently, Rbfox2 was 1.5-fold enriched (right side). **f)** Overexpression of miR-92a-3p in N2A cells led to >2-fold enrichment of miR-92a-3p (left side) and subsequently to a decreased abundance of Rbfox2 (right side). **g)** Overexpression of mir-92a-3p in N2A cells reduced the RBFOX2 protein level significantly. **h)** Representative image of the western blot for RBFOX2 levels following miR-92a-3p overexpression in N2A cells. **i-j)** miR-92a-3p induces splicing changes by regulating Rbfox2 expression. Overexpression of miR-92a-3p induced exon skipping in Lrp8 (i) and exon inclusion in Dnm2 (j). MiR-92a-3p-OE and Rbfox2-OE was performed in N2A cells and in the cerebral cortex, respectively. Data are shown as mean ± standard error of the mean. *p<0.05, ** p<0.01, *** p<0.001, ****p<0.0001, Wald z-test from DESeq2 in b,c; paired t-test or paired Wilcoxon test in d-g, linear mixed effects analysis in i-j.

Therefore, we focused our attention on PTBP2, which is a high-confidence protein-protein interaction partner of RBFOX2 (Supplementary Figure S7b). To investigate whether PTBP2 and RBFOX2 bind to overlapping splicing targets, we screened for PTBP-motifs upstream and RBFOX-motifs downstream of alternatively spliced exons and for direct RNA-protein binding sites by reanalyzing iClip data^38,58^. Roughly 45% of the significant inclusion events (320) had the combination of RBFOX-motif downstream with a PTBP-motif upstream. The majority of these were also highly conserved (Supplementary Figure S7a). Most included exons (547) harbored either a downstream RBFOX-motif and/or an RBFOX-binding site and simultaneously an upstream PTBP-motif or a PTBP-binding site (Figure 5c).

To corroborate the antagonistic splicing activity of RBFOX2 and PTBP2, we re-analyzed wildtype and *Ptbp2*-KO data of P1 mouse neocortices^58^. We detected 707 alternatively spliced exons in *Ptbp2*-KO (429 included, 187 excluded), with 128 events shared with *Rbfox2*-OE. This represented a 5.24-fold enrichment compared to the expected overlap (p < 0.0001, Supplementary Figure S8a). Supporting the proposed antagonistic relationship between both RBPs, most shared target exons were spliced in the same direction upon *Ptbp2*-KO and *Rbfox2-OE* (85.2%, Supplementary Figure 8c). Furthermore, we observed a moderate positive correlation between exon inclusion levels in *Ptbp2*-KO and *Rbfox2-*OE samples (R = 0.2993, p < 0.001, Supplementary Figure 8b). Next, we analyzed the splicing pattern in our data of 230 PTBP2-repressed exons reported to be relevant specifically during neocortical development^58^. *Rbfox2*-OE induced alternative splicing in 67 of these PTBP2 targets. Importantly, 55 of the significant events showed exon inclusion in the *Rbfox2*-OE group (Figure 5d, Supplementary Data 6). Among the antagonistically spliced genes was *Ncam1* with an abnormal early inclusion of exon18 upon *Rbfox2-*OE (chr9:49,418,152-49,418,953, Supplementary Figure S6c) which leads to the production the NCAM180 isoform. Interestingly, NCAM140 together with NCAM180 are the predominant isoforms for neurons, whereas NPCs express NCAM120 and NCAM140^59^.

Another PTBP2-excluded^58^ and RBFOX2-included cassette exon event was identified in *Kif21b* (chr1:136,101,209-136,101,389, Supplementary Figure S6d), which is an important regulator of microtubule dynamics.

Furthermore, exonN (chrX:73,284,421-73,284,478) of the *Flna* transcript was significantly included upon *Rbfox2-*OE (Supplementary Figure S6e). ExonN was also specifically repressed in NPCs and included in neurons, while exon inclusion was strongly elevated from E14.5 to E16.5. The iClip^38,58^ data also confirmed protein binding for RBFOX2, PTBP1, and PTBP2 in flanking sequences of exonN. *Flna* is an important regulator of neuronal migration^60^, and its splicing by PTBP1 in NPCs is associated with shaping the NPC-to-neuron transition during cortical development^8,58^. To validate the biological relevance of the proposed antagonistic splicing regulation in the context of brain development, we next performed an in-depth cell-type-specific characterization of the expression of *Rbfox2* and *Ptbp2* in the embryonic neocortex using the scRNA-seq data by Ruan and colleagues^32^. Force-directed graph embedding and PAGA visualization allowed us to reconstruct the differentiation trajectories from NECs to different types of cortical neurons and glial cells (Figure 5e). While *Ptbp2* was expressed across a broad range of cell types, its levels were highest in NECs, RGCs, and mitotic IPCs, where *Rbfox2* was mostly absent (Figure 5f). To elucidate regulation in the neuronal lineage, we next examined expression patterns of both proteins along differentiation pseudotime (Supplementary Figure S7d), specifically in the NECs-RGCs-IPC-immature neurons-layer V-VI neuron trajectory (Figure 5g). This analysis unequivocally identified late IPCs and early immature neurons as the turning point from *Ptbp2*-dominated to higher *Rbfox2* expression, correlating with the transition from progenitor-like to neuronal splicing patterns.

Notably, while the percentage of cells uniquely expressing *Ptbp2* decreased along the differentiation trajectory and the opposite effect was present for *Rbfox2,* a fraction of cells co-expressed both genes in progenitor and differentiated cells (Figure 5h). Therefore, these results support the notion that splicing regulation is not dictated solely by the presence or absence of either protein, but rather by their relative expression levels, thus lending credibility to our hypothesis that premature overexpression of *Rbfox2* in our study interfered with the splicing activity of PTBP2, potentially leading to the impaired differentiation and migration phenotype we observed.

### miR-92a-3p regulates the expression of *Rbfox*2 and impacts its splicing activity

To investigate whether other PTBP proteins could be relevant in the transition from neural progenitors to differentiated neurons, we next characterized their developmental-stage and lineage-specific expression (Figure 6a, Supplementary Figure S9e). While *Ptbp3* was restricted to the glial lineage, *Ptbp1* was present at low levels in undifferentiated cells and at earlier developmental stages in accordance with previous reports^39,58,61^. These characteristic expression patterns imply strict transcriptional regulation Notably, *Ptbp1* repression is mediated via miR-124-3p, which is a highly specific neuronal miRNA (Figure 6b)^17,62^. Furthermore, miR-124-3p becomes significantly up-regulated in the transition between E14 and E17 of cortical development (Figure 6b)^62^.

Therefore, we hypothesized that miRNAs could also be important regulators of *Rbfox2* expression. In a previous study, we identified 36 miRNAs that were expressed in the embryonic cerebral cortex and predicted to target the 3’ untranslated region (3’ UTR) of *Rbfox2.* By further inspecting these miRNAs, we observed that most of them were upregulated at E14, followed by a significant reduction of expression levels at the subsequent developmental stages E17 and P0 (Supplementary Figure S9a)^62^. This prompted us to hypothesize that miRNAs might mediate the low expression levels of *Rbfox2* in NPCs. Specifically, miR-92a-3p was identified as a hub microRNA at E14 during cortical development^62^, and it has a predicted binding site in the 3’ UTR of *Rbfox2* which is highly conserved among vertebrates. Furthermore, the expression of miR-92a-3p was higher in NPCs compared to neurons and decreased gradually from E14 to P0 (Figure 6c). Together, this data suggests that miRNA-92a-3p might contribute to silencing *Rbfox2* in NPCs at early developmental stages.

To test this hypothesis, we performed luciferase assays using a fragment of the 3’ UTR of *Rbfox2,* which contained the miR-92a-3p binding site downstream of the *Renilla* luciferase gene. Co-transfection of this reporter plasmid into HEK293 cells together with a miR-92a-3p mimic or a control revealed a reduced luciferase activity of more than two-fold in cells transfected with miR-92a-3p compared to the control. This finding confirmed the gene silencing effect of miR-92a-3p on *Rbfox2* (p < 0.01, Figure 6d). To investigate the mir-92a-3p-mediated suppression of *Rbfox2 in vivo,* we electroplated mouse embryos at E14.5 with a miR-92a-3p-silencing construct. Expression of miR-92a-3p was reduced more than 10-fold compared to control samples (p < 0.0001, Figure 6e). Following miR-92a-3p reduction, we observed a significantly increased *Rbfox2* expression at E16.5 in the miR-92a knock-down samples compared to the control (p < 0.01, fold change = 1.571; Figure 6e). To further corroborate these results, we overexpressed miR-92a-3p in an N2A cell culture. This resulted in a significant increase of miR-92a-3p abundance, as measured by TaqMan assay (p < 0.05, fold change = 2.442; Figure 6f). Importantly, both the mRNA expression (p < 0.05, fold change = 0.748 relative to control; Figure 6f) and the protein levels of RBFOX2 (p < 0.05, fold change = 0.707 relative to control; Figure 6g,h) were reduced significantly upon miR-92a-3p-overexpression.

To investigate if miR-92a-3p-mediated silencing of RBFOX2 also impacts its alternative splicing activity, we performed RT PCR analysis of target exons in five genes that we previously detected as alternatively spliced upon *Rbfox2* overexpression in the embryonic cerebral cortex. We employed long-read Nanopore sequencing of PCR amplicons to quantify exon inclusion levels in miR-92a-3p-OE and control samples. We observed only exon skipping for three of the target genes in both experimental groups – *Clasp2*, *Tead1, and Dlg2* (Supplementary Fig. S9b-d).

However, inclusion levels for the target exon in the *Lrp8* gene were significantly lower in the miR-92a-3p-OE compared to control samples. In contrast, *Rbfox2-*OE was associated with higher exon inclusion in the embryonic cerebral cortex (Figure 6i), whereas we previously observed exon skipping upon *Rbfox2-*OE. Similar regulatory effects were observed for *Dnm2*, where *Rbfox2*-OE significantly increased exon inclusion levels compared to control, while miR-92a-3p-OE showed the opposite effect with significantly reduced exon inclusion (Figure 6j). These results therefore serve as proof of principle that miR-92a-3p influences expression levels of RBFOX2, which in turn also impacts the splicing outcome of RBFOX2 targets.

## Discussion

RBFOX2 is a crucial component of the complex regulatory network that orchestrates cerebral cortical development. We revealed that misexpression leads to a global disruption of the alternative splicing landscape with a significant overrepresentation of high-confidence NDD risk genes (Supplementary Figure S4). Consequently, aberrant RBFOX2 splicing activities could produce target gene isoforms that lead to impaired brain development and disorders of the central nervous system in humans^63,64^. For instance, a mutation in *RBFOX2* has been detected previously in a patient with hypoplastic left heart syndrome and NDD^21^. Surprisingly, however, limited research has been conducted on the clinical significance of RBFOX2, as most studies focus on RBFOX1. Mutations in *Rbfox1* have been consistently linked with autism spectrum disorder^65–68^ and susceptibility to epilepsy^69–71^. Interestingly, *Rbfox1* knock-out resulted in higher seizure susceptibility^72^, whereas overexpression was also linked with epilepsy and cortical malformation^73^. Furthermore, reduced cytoplasmic RBFOX1 levels in the prefrontal cortex of individuals with schizophrenia^24^ decrease the stability of *Vamp1*, a schizophrenia-associated gene, in PV^+^ interneurons^25^. This reduction impairs inhibitory drive from PV^+^ interneurons, contributing to schizophrenia pathogenesis^74^.

Though RBFOX1 and RBFOX2 have largely overlapping alternative splicing targets^35^, the divergent expression patterns we observed here and in previous research indicate that both proteins have common but also distinct functions^23,72,75,76^. To disentangle the specific role of RBFOX2 in the developing brain, we employed an overexpression model instead of a knock-down strategy, as the members of the RBFOX family can compensate for the absence of one of their paralogues^23^. Interestingly, early *Rbfox2*-OE caused a premature detachment and migration of neural cells to the lIZ but not further (Figure 4a). This was accompanied by significant splicing events in key components of the Reelin pathway^48,49^ , which plays a crucial role in the migration and localization of cortical neurons and Purkinje cells in the cerebellum. Gehman and colleagues previously showed that the Reelin-binding *Lrp8* gene was alternatively spliced upon *Rbfox2* knock-out. Consequently, Purkinje cells failed to detach and therefore remained near their origin in the VZ^23^. Together with our results, this indicates an important role of RBFOX2 in the initiation and control of cell migration in different parts of the developing nervous system.

Apart from the altered migration, we also observed impaired neuronal differentiation in *Rbfox*2-overexpressing cells (Figure 4b-d). RBFOX2 was specifically upregulated in the transition from NPCs to newly differentiated neurons (Figure 1e), highlighting its role in neural differentiation. Accordingly, we detected a strong alternative splicing pattern in *Numb,* which is a key player in asymmetrical neural stem cell division, determining the cell fate of progenitor cells and neurons^77^. Different isoforms of *Numb* can either promote neuronal differentiation or enhance cell proliferation^78^, giving a possible explanation for the observed phenotype of decreased neuronal differentiation in our study.

Interestingly, *Rbfox2* expression negatively correlates with expression patterns of the PTBP family. In particular, PTBP1 is a key regulator of non-neuronal splicing in progenitor cells and its repression is required to induce the transition to neurons^17^. In contrast, PTBP2, which is reported to be a weaker repressor of alternatively spliced exons than PTBP1^40,79^, is down-regulated by PTBP1 via poison exon inclusion leading to nonsense-mediated decay^17,38,61,80^. Remarkably, our data provided strong hints for an antagonistic splicing relationship between RBFOX2 and PTBP. Specifically, RBFOX2 and PTBP2 shared a set of targets that undergo alternative splicing during cerebral cortical development (Figure 5d). Although PTBP2 typically promotes the exclusion of these antagonistically spliced exons, RBFOX2 induced the opposite effect of exon inclusion. Therefore, the premature overexpression of RBFOX2 in our experimental set-up possibly interfered with the splicing program of PTBP2 which could explain the aberrant cellular phenotype and altered splicing landscape we observed. Indeed, several targets that are spliced antagonistically between RBFOX2 and PTBP2 play crucial roles, for example, in regulating neuronal differentiation (*Flna*)^8^, neurite outgrowth (*Ncam1*), and microtubule dynamics (*Kif21b*). Furthermore, PTBP-recognized motifs are reportedly enriched upstream of alternatively spliced exons that show high inclusion levels in the brain, heart, and skeletal muscles^53^. These tissues are known expression sites of RBFOX2^81^ with a concurrent low expression of PTBP1^53^. Therefore, our results and existing reports point to a complex antagonistic interplay between these proteins in regulating neuronal differentiation^4,8^. Supporting the concept of such dose-sensitive antagonistic interactions among RNA-binding proteins, Ellis and colleagues recently reported that even a modest increase in RBFOX expression levels was sufficient to alter splicing events dependent on another RBP, MBNL1^82^. Furthermore, RBFOX-concentration-dependent splicing regulation has been described in relation to binding to secondary motifs. Accordingly, high RBFOX2 protein levels during neuronal differentiation can mediate exon inclusion through binding to low-affinity motifs^83^. These findings illustrate how nuanced temporal and spatial RBP expression levels enable cell- and developmental-stage-specific gene regulation during brain development.

Indeed, several post-transcriptional regulatory mechanisms controlling RBFOX levels have been reported already. For instance, sno-lncRNAs can act as sequestering sites for excess RBFOX2 proteins^75^. Furthermore, RBFOX3 can splice poison exons into *Rbfox2* leading to NMD^84^. AnotherI potential layer of regulation is a putative *Rbfox1-* and *Rbfox2-*specific feedback loop. Accordingly, we observed significantly reduced levels of *Rbfox1* following the overexpression of *Rbfox2* (Figure 2e). Similarly, a knock-out of *Rbfox1* was previously reported to lead to an increased level of RBFOX2^72^ and vice versa^23^. Furthermore, both RBFOX1 and RBFOX2 contain RBFOX binding sites near a highly conserved 93-nucleotide exon encoding the second half of the RRM. RBFOX binding leads to exon skipping, thereby resulting in a RbfoxΔRRM isoform with significantly reduced RNA binding affinity^85^. In line with this, the *Rbfox2* overexpression in early-stage cortical cells in our study did not result in a complete splicing switch with all regulated exons being either fully spliced in or spliced out.

miRNA binding to 3’ UTRs represents another key post-transcriptional regulatory mechanism for controlling gene expression and protein levels. In a study investigating the expression patterns of miRNAs in the embryonic cerebral cortex, we recently highlighted targeting of RNA binding proteins as one of the major processes through which miRNAs impact brain development^62^. Similarly to the reported silencing of *Ptbp1* in neurons via miR-124-3p^17^, here we showed for the first time that the NPC hub miRNA 92a-3p suppresses the expression of *Rbfox2.* Conversely, silencing miR-92a-3p in the embryonic neocortex led to a significant *Rbfox2* upregulation. In accordance with our results, miR-92a-3p was previously reported to be enriched in intermediate progenitor cells in humans^86,87^ and, interestingly, differentially expressed in neurodevelopmental and mental disorders. For instance, levels were elevated in blood^88^ and serum^89^ of patients with schizophrenia. In contrast, miR-92a-3p was reported to be downregulated in individuals with autism^90,91^. Given the overlapping splicing targets between RBFOX2 and its paralogue RBFOX1, whose mutations have been consistently implicated in such disorders, disruption of the miR-92a-3p/RBFOX2 axis represents an intriguing mechanism for the emergence of diseases of the central nervous system as well as a potential therapeutic target. In particular, future research should focus on elucidating the link between altered miR-92a-3p expression levels and aberrant splicing activities that contribute to disease pathogenesis.

Collectively, our results underscore the crucial role of RBFOX2 in the developing cerebral cortex and shed light on the complex interplay with the PTBP family in antagonistically regulating alternative splicing of common target genes. Additionally, we show for the first time that low RBFXO2 expression levels in neural progenitors are mediated via miR-92a-3p, thereby highlighting miRNAs as key components of the intricate regulatory network that shapes embryonic brain development.

## Methods

### Animals

Hes5::GFP mice have been described previously^92^. For *in utero* electroporation, timed mated c57BL6/JRj female mice were purchased on gestation day 12 from Janvier (Janvier Labs, Le Genest-Saint-Isle, France), a certified international breeder. Mice were housed in type II long filter top cages (Tecniplast, Buguggiate, Italy) and kept in a 12 h light/dark cycle in a temperature and humidity-controlled animal room (22 ± 2°C, 55 ± 5%). Water and food (ssniff M-Z Extrudat, ssniff, Soest, Germany) were supplied *ad libitum*. Mice were treated according to the protocols approved by the local authorities (Landesuntersuchungsamt Rheinland-Pfalz, license number 23 177-07/G 13-1-089).

### Constructs

HA-tagged full-length *Rbfox2* was cloned into the XhoI and BglII sites of the pCAGGS-mCherry vector using the following primer combinations: *Rbfox2* 1A full-length forward 5′- actgctcgaggccaccatgtacccatacgatgttccagattacgctgcggaaggcggccaggcg-3′, with *Rbfox2* reverse 5′-actgagatcttcacgtcacttcagtagg-3′. The miR-92a pre-miRNA including approximately 150 bp of flanking genomic sequence was cloned into the BamHI and EcoRI sites of the pcDNA3.1+ vector using the following primer combinations: miR-92 forward 5′-agtcggatcctttagcgttggaaagtggcc-3′ with miR-92a reverse 5′-agtcgaattctaagttgaggtgtgggtggg-3′. A fragment of the *Rbfox2* 3’UTR containing the miR-92a binding site was cloned into the XbaI site of the pGL3P vector using the following 5’-end phosphorylated oligonucleotides: Sense [Phos] 5′- CTAGACCCCAGTTCATGAGGCCTGGCTATTGCAATATTTACTAGTAGAGGACTCTA TAGCT-3′, Antisense [Phos] 5′-CTAGAGCTATAGAGTCCTCTACTAGTAAATATTGCAATAGCCAGGCCTCATGAACTGGGGT-3′. Sense and Antisense oligos were annealed in annealing buffer (10mM Tris-HCL; pH 7.5; 0.1 M NaCl; 1 mM EDTA) and cloned into the XbaI site of the pGL3P vector.

### Cell Culture, transfections, and luciferase assays

N2A and HEK293 cells were cultured in Dulbeccós Modified Eagle Medium (DMEM) with 10% fetal bovine serum as well as 1% Penicillin/Streptomycin. For Luciferase assays, HEK293 cells were seeded on 12-well plates at a density of 90,000 cells per well. Twenty-four hours later, the cells were transfected with 400 ng of the constructs cloned into the pGL3P vector, 600 ng miRNA construct and 10 ng pRL-TK vector using the calcium phosphate method. After 48 hours, cells were lysed, and luciferase activity was measured with a CentroXS LB 960 luminometer (Berthold Technologies, Germany). N2A cells were transfected with Transfectin reagent (Biorad, CA, USA). The day before the experiment, for transfection, N2A cells were seeded at a density of 250,000 cells per well on 6-well plates. Twenty-four hours later, the cells were transfected with 4000 ng miRNA construct per well. The cells were lysed and processed for RT-qPCR, Western blot and Nanopore sequencing analysis 48 hours after transfection.

### RT-qPCR and Western blot

Small RNAs were isolated using the miRNeasy Kit (Qiagen, Germany), and total RNA was isolated using the High Pure RNA isolation kit (Roche, Switzerland). TaqMan MicroRNA Assays (Applied Biosystems, CA, USA) were used for miRNA RT-qPCR experiments. The miRNA quantities are relative to U6 snRNA. Total RNA was reverse transcribed using the RevertAid First Strand cDNA Synthesis Kit (Fermentas, Sankt Leon Rot, Germany). For qPCR, SYBR Green Mix was used (TB Green Premix Ex Taq II (RR82WR, Takara, Japan)).

For Western blot analysis, the cells were lysed, and the protein lysates were generated from cell pellets using Magic Mix (48% urea, 15mM Tris pH7.5, 8.7% Glycerin, 1% SDS, 143mM β-mercaptoethanol) containing protease and phosphatase inhibitors (cOmplete Tablets easypack, PhosSTOP easypack, Roche) and transferred to a QIAshredder column (Qiagen). After centrifugation at 13500g for 2 minutes, the solution was transferred into a fresh tube and frozen at -80°C until needed. Later, the proteins were resolved by electrophoresis on SDS acrylamide gels. After transferring the proteins to PVDF membranes, Western blot analyses were performed with antibodies specific for RBFOX2 (rabbit-Anti Rbm9; Bethyl Laboratories, TX, USA) and GAPDH (mouse-Anti GAPDH ab8245, Abcam, UK). Quantification of the bands was performed with ImageJ and normalized to GAPDH band intensity.

### *In utero* Electroporation

For *in utero* electroporation, embryonic day E13.5 pregnant female mice were deeply anesthetized with isoflurane, and the uterine horns were exposed. For overexpression, control or *Rbfox2* expression vectors (4 µg/µl) were injected with either pCAG-mCherry or pCAG-eGFP (0.5 µg/µl) into the lateral ventricle of each embryo. For electroporation 5 pulses of 40 V for a pulse length of 50 ms at 950-ms intervals were applied (Nepagene, Japan). Subsequently, the uterine horns were returned to the abdominal cavity. After 48 h, the mice were sacrificed by cervical dislocation. The embryos were collected and sacrificed by decapitation. For immunostaining experiments, the brains were isolated, fixed in 4% paraformaldehyde, cryoprotected in 15% and 30% Sucrose solutions, embedded in Cryomatrix (Thermo Scientific, MA, USA), and cryosectioned into 20 µm-thick brain slices.

For RNA-sequencing, the brain’s GFP^+^ area was dissected and digested with trypsin. After stopping the digestion by adding 20% FBS in DMEM and washing with PBS, cells were resuspended in PBS and passed through a 100 µm filter to achieve a single-cell suspension. GFP^+^ cells were then collected by fluorescent activated cell sorting (FACS).

### Immunostaining and Imaging

For immunostaining, brain cryosections were fixed in 4% paraformaldehyde (PFA) at room temperature (RT) for 10 minutes. After two washes with PBS, antigen retrieval was performed by incubating the slides in 10 mM sodium citrate with 0.5% Tween-20, pH 6, at 80°C for 20 minutes. Subsequently, the sections were blocked in PBS with 0.2% Tx-100 containing 2% sheep serum. The sections were then incubated with primary antibody (rabbit anti-RBM9 (Bethyl, TX, USA), mouse anti-RBFOX1 (MABE985, Merck/Millipore, Germany/MA, USA, Burlington, MA, USA), rabbit anti-beta III Tubulin (ab18207, Abcam, UK), rabbit Anti-PAX6 (901301, Biolegends, CA, USA), chicken Anti TBR2 (ab34735, Millipore, MA, USA), rabbit Anti-SATB2 (ab34735, Abcam, UK)) in blocking solution overnight at 4°C. The sections were washed three times in PBS 0.2% Tx-100 and incubated in blocking solution containing Alexa-488- or Alexa-594-conjugated secondary antibody (Invitrogen, CA, USA) for 3 hours at RT. After three washes in PBS, the sections were embedded in Fluoromount-G with DAPI (00-4959-52, Invitrogen).

The sections were viewed using a laser-scanning confocal microscope (LSM710), and images were acquired with the zen software (Zeiss, Germany). The images were analyzed, and the cells were counted using ImageJ software^93^.

### RNA-sequencing

NGS library prep was performed with Clontech SMARTer Ultra Low Input RNA Kit (Clontech, CA, USA) for cDNA generation, followed by NuGEN Ovation Ultralow v2 System (NuGEN, CA, USA) for library preparation. Libraries were profiled in a DNA 1000 chip on a 2100 Bioanalyzer (Agilent Technologies, CA, USA) and quantified using the Qubit dsDNA HS Assay Kit, in a Qubit 4.0 Fluorometer (Invitrogen by Thermo Fisher Scientific, MA, USA). All six samples were pooled in equimolar ratio and sequenced on an Illumina NextSeq 500/550 device (Illumina, umina, CA, USA).

### RNA-sequencing analysis

The paired-end reads were trimmed using BBDuk (version 39.01)^94^. The trimmed reads were mapped to the Gencode reference genome of Mus musculus mm39 (released 19.10.2022) using STAR (version 2.7.10b)^95^ and the count per gene and sample was determined using FeatureCounts provided by SubRead (version 2.0.6)^96^. We used DESeq2 (version 1.40.1)^97^ in R (version 4.3.0)^98^ to analyze differentially expressed genes with default settings. Since *Rbfox2* was heavily overexpressed, it was removed from the volcano plot for improved visualization. Genes with an adjusted p-value (Benjamini-Hochberg method) < 0.05 were considered to be differentially expressed.

rMATS (version 4.1.2)^99^ with default settings was used to detect alternatively spliced genes between the *Rbfox2*-OE and control samples. We did not include novel splice events in our analysis. A splicing event with a false discovery rate (FDR) adjusted p-value < 0.01 and an inclusion level difference |ΔPSI| > 0.1 was considered significant. Data processing was done with numpy (version 1.23.5)^100^ and pandas (version 2.0.1)^101^ in Python (version 3.10.12)^102^.

Plots were generated if not specified otherwise with Matplotlib (version 3.7.1)^103^ and seaborn (version 0.12.2)^104^.

The overlap between significantly alternatively spliced genes and significantly differentially expressed genes was visualized in an upset plot (version 0.9.0)^105^. Sashimi plots were created using rmats2sashimiplot within a Docker container using the image xinglab/rmats2sashimiplot. Due to its MISO backend, PSI values may differ from rMATS results.

### Nanopore Amplicon Library Preparation and Sequencing

Amplicon libraries were prepared using the PCR Barcoding Expansion kit (EXP-PBC001) in combination with the Ligation Sequencing Kit V14 (SQK-LSK114) from Oxford Nanopore Technologies (Oxford Nanopore Technologies, UK). Briefly, an initial PCR amplification for specific RBFOX2 targets was performed on cDNA samples using Herculase II Fusion DNA Polymerase (Agilent Technologies, CA, USA) with gene-specific primers tailed at the 5′ end. The forward primers were appended with 5′-tttctgttggtgctgatattgc-3′ and the reverse primers with 5′- acttgcctgtcgctctatcttc-3′. PCR products for each sample were pooled equimolarly based on volume. This was followed by a second round of PCR to incorporate sample-specific barcodes. After amplification, each barcoded sample was cleaned using AMPure XP beads (Beckman Coulter, CA, USA), quantified, and pooled in equimolar amounts. The pooled library was subjected to end-repair and dA-tailing, followed by ligation of sequencing adapters using the Ligation Sequencing Kit. Adapter-ligated libraries were cleaned with AMPure XP beads (Beckman Coulter, CA, USA) and stored at 4°C for short-term use. Prior to sequencing, the PromethION flow cell was primed and the prepared library was loaded following the manufacturer’s protocol.

### Nanopore RNA-sequencing analysis

Base calling of raw pod5 files was performed using Dorado (version 1.0.2, Oxford Nanopore Technologies, UK), and reads were aligned to the Gencode reference genome of Mus musculus mm39 using samtools (version 1.16.1) and minimap2 (version 2.26-r1175). For each amplified target region, genomic coordinates for the upstream exon, target exon, and downstream exon were extracted from the rMATS output of the *Rbfox2*-OE analysis. Sequencing reads mapping to these defined regions were extracted using pysam (version 0.21.0) and subjected to quality filtering with a minimum mapping quality threshold of 10. Read classification was based on exon mapping patterns to quantify alternative splicing events. Reads were categorized as supporting exon skipping when they aligned to both upstream and downstream exons while failing to map to the target exon. Conversely, reads mapping to all three exons (upstream, target, and downstream) were classified as supporting exon inclusion. Reads that did not meet either classification criteria were excluded from downstream analysis. Statistical analysis of exon inclusion levels between control and miR-92a-3 overexpression samples was performed using linear mixed effects models with the lme4 R package (version 1.1.37) followed by an analysis of deviance implemented in the car package (version 3.1.3). Experimental group and miR-92a-3p overexpression fold change were included as fixed effect in the model and sample replicates as random effects.

### Analysis of publicly available transcriptomic data sets (bulk RNA-seq)

To further characterize the alternative splicing landscape during cortical development, we re-analyzed two publicly available data sets. The study by Weyn-Vanhentenryck et al. contains RNA-sequencing of the mouse cortex from nine time points (E14.5, E16.5, P0, P4, P30, 4 months, and 21 months). Sequencing files were downloaded from the NCBI Short Read Archive, accession number SRP055008. The study by Liu et al. contains RNA-sequencing from NPCs and neurons purified from mouse cortices at E15.5, sequencing files were obtained from the Gene Expression Omnibus, accession number GSE96950. Furthermore, we re-analyzed transcriptomic data from *Ptbp2* knockout mouse cortices at P1, obtained from the Gene Expression Omnibus (accession number GSE84803)^58^, and triple knockout (*Rbfox*1/2/3) mouse embryonic stem cells, retrieved from the Short Read Archive (accession number SRP128054)^33^. The sequencing data were processed according to the protocol specified under the section “RNA-sequencing analysis”. The developmental data from Weyn-Vanhentenryck et al. did not allow a pairwise statistical test with rMATS, therefore we conducted a pairwise Fisher’s exact test in scipy (version 1.10.1)^106^ for consecutive time points and corrected p-values for multiple comparisons by using Benjamini Hochberg’s procedure. An event was considered significantly developmentally alternatively spliced when two successive time points had an adjusted p-value < 0.01 and |ΔPSI| > 0.1.

We used a Monte Carlo simulation to statistically assess the overlap of alternative splicing events (cassette exons) between *Rbfox2*-OE and either t*Rbfox*-KO^33^ or *Ptbp2*-KO^58^ conditions (see Monte Carlo simulation for details).

### Analysis of publicly available transcriptomic data sets (scRNA)

We obtained a single-cell RNA sequencing dataset for the developing mouse neocortex (GSE161690) generated by Ruan and colleagues^32^ at developmental stages E10.5, E12.5, E14.5, E15.5, E16.5, and E18.5. The original cell type annotations provided by the authors were retained for subsequent analysis. Data processing and analysis were performed using scanpy (version 1.11.3)^107^. Initial data processing involved normalizing the gene expression counts for each cell to a target sum of 1,000, followed by a log10(x+1) transformation. We selected the 500 highly variable genes using scanpy.pp.highly_variable_genes. Principal Component Analysis was then performed using the top 16 principal components. Subsequently, a neighborhood graph was computed with 10 neighbors per cell. This neighborhood graph served as the basis for generating the UMAP for visualization.

To infer the developmental trajectories between different cell types, we applied the partition-based graph abstraction (PAGA) routine. The root for pseudotime calculation was set to the “Neuroepithelium-a” cell type. Our specific interest was in the differentiation pathway of progenitor cells into neurons. Therefore, we focused our analysis on the inferred trajectory path encompassing “Neuroepithelium-a,” “Neuroepithelium-b,” “RGC-a,” “IPC-a,” “immature neuron,” and “Layer V-VI-a” cell types. Along this specific trajectory, we extracted the expression values for *Rbfox2* and *Ptbp2* genes. To model the expression trends of these genes over pseudotime, we performed spline interpolation using scipy (version 1.10.1)^106^. We performed cubic spline interpolation using scipy.interpolate. UnivariateSpline to model gene expression trends over cell type in order of the trajectory and along pseudotime. To estimate the 95% confidence interval of these fits, we applied a non-parametric bootstrapping approach with 1,000 iterations, resampling pseudotime-expression pairs and calculating percentiles of the bootstrapped spline predictions.

### Analysis of publicly available iClip data sets

To validate which significant splicing events could be attributable to direct RBFOX2 binding, we re-analyzed publicly available iClip data for RBFOX2 expressed in V6.5 mESCs. The data was downloaded from NCBI Gene Expression Omnibus, accession number GSE54794^38^. This dataset was supplemented by iClip data for PTBP1 and PTBP2, downloaded from the NCBI short read archive, accession number SRP080878^58^.

The iClip data was analyzed in several steps. First, we extracted the unique molecular identifiers from the reads using UMI tools (version 1.1.4)^108^ and trimmed the adapters with BBDuk (version 39.01)^94^. The trimmed reads were aligned to the reference genome mm39 from Gencode (released 19.10.2022) using STAR (version 2.7.10b)^95^. Following the alignment, we removed PCR duplicates by leveraging the unique molecular identifiers with UMI tools. Crosslinking sites were identified as peaks using Clipper^109^, which was run within a Docker container based on the image brianyee/clipper:61d5456.

### Inferring the alternative splicing trajectory during cortical development

To analyze the effect of *Rbfox2*-overexpression on the alternative splicing pattern concerning normal cortical development, we used the SplicePCA tool^42^ integrating our data set with the developmental data of Weyn-Vanhentenryck et al.^4^ and Liu et al.^29^. We selected events that did not have missing data for any sample and were significantly alternatively spliced in our data and the publicly available data. The developmental trajectory was fitted using spline regression with bootstrap resampling (n=500) to generate 95% confidence intervals. To investigate how splicing events are correlated with the principal components, we then computed the loading matrix A

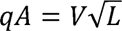

where V is the matrix of eigenvectors, L is a diagonal matrix with the corresponding eigenvalues and loadings correspond to correlations between the principal components and the original variables^110^.

For each principal component, we then performed a gene set enrichment analysis (GSEA) using the clusterProfiler package (version 4.8.1)^111,112^ in R v4.2.2 with the biological process ontology, miGSSize of 100, maxGSSize of 500, and an adjusted p-value cut-off of 0.05. The loadings for each gene on the respective principal component were used as input for the GSEA.

### Inferring the direction of exon inclusion

To assess whether *Rbfox2-*OE is related to a mature (neuronal) or immature (NPC-specific) splicing pattern, we leveraged the NPC and neuron RNA-sequencing data from Liu et al.^47^. By intersecting our significant alternative splicing results with the Liu et al. data set, we assigned the *Rbfox2*-OE splicing events to three categories: 1) Neuronal direction when the exon inclusion level of the respective gene was significantly higher in neurons than in NPCs in the Liu et al. data; 2) NPC direction when the inclusion level was significantly lower in neurons than in NPC; 3) non-regulated, when the inclusion level was not significantly altered, or the splicing event was not detected in the Liu et al. data set. The results of this analysis were visualized with a chord diagram using the circlize package (version 0.4.15)^113^ in R.

### Motif enrichment analysis

We performed a *de novo* motif enrichment analysis to identify enriched motifs in the intronic regions located upstream and downstream of alternatively spliced exons. The intronic sequences around alternatively spliced exons (enriched and suppressed exons separately) and a set of background sequences that were not alternatively spliced (FDR > 0.8 and |ΔPSI| < 0.1) were loaded from the mm39 reference genome (Gencode released 19.10.2022) using pysam (version 0.21)^114^.

All possible pentameric sequences and their occurrence in the up- and downstream intronic sequences were counted in a sliding window of 50 nt. For each motif, we then subtracted its occurrence in the respective background sequences. Next, motif frequencies were transformed to z-scores using scipy. A motif was considered to be enriched when it had a z-score > 4 (p < 0.0001, two-sided z-test).

The iClip peaks were used to validate these predicted motifs. We again screened for the occurrence of every possible pentameric sequence within the peaks of RBFOX2 and the control, subsequently subtracted the respective values, and performed z-scoring. Next, the z-scores of the iClip and RNA-Sequencing motif enrichment analysis were correlated using scipy.

### Motif co-occurrence analysis

The co-occurrence of motifs was analyzed similarly to the motif enrichment analysis. 400 nt of intronic sequence flanking the alternatively spliced exon or the exon itself was loaded with pysam (version 0.21.0)^114^ and screened for the presence of a PTBP-associated CU-rich motifs (UCUCU, UUUCU, UCUUU, CUCUC, UCUCC, CUCUU, UUCCU)^115^ with a z-score > 4 upstream and RBFOX2 canonical motif downstream (GCAUG, UGCAU, CAUGC). Moreover, we integrated confirmed binding sites from the iClip data for RBFOX2^38^, PTBP1, and PTBP2^58^. We visualized this using an upset plot and only included overlapping sets with a minimum size of 1.

Further, we examined evolutionarily conserved motif occurrences around these exons by filtering for motifs with a phastcons 35-way^116^ score > 0.8.

### Motif localization map

Next, we mapped the localization of the enriched motifs identified in the previous analysis on the alternatively spliced exon and the flanking exons. To the end, we computed the motif density in a sliding window of 50 bp for all up- and downregulated events and background (non-regulated events). Motif density corresponds to the fraction of the 50 nt sliding window that is covered by the motif. The estimates were corrected by subtracting the background density for each motif. This analysis was limited to 150 nt of exonic and 400 nt of intronic sequences spanning up- and downstream from the intron/exon boundaries.

### STRING network analysis

Potential interaction partners of RBFOX2 were identified with STRING using all potential interaction sources (version 12.0 beta)^117^ allowing 20 interacting proteins and showing confidence as edges. Interactions to RBFOX2 with a confidence score higher than 0.7 were considered high confidence.

### Statistical analysis of immunohistochemistry data

Two groups were compared statistically using a t-test or a Wilcoxon test when assumptions of the parametric test were violated, and a data transformation did not improve the model fit. More than two groups were analyzed using one-way or two-way analysis of variance followed by Tukey’s post hoc test or post hoc comparison of model means. Data were transformed using a log_e_ or an arcsine square root transformation when assumptions of normality and homogeneity of variance were violated. All p-values are two-tailed and a p-value < 0.05 was considered statistically significant. Statistical analysis was performed using R v4.2.2 or Python (version 3.10.12)^102^ using scipy (version 1.10.1)^106^.

### Overlap of alternatively spliced genes with NDD risk genes

Alternatively spliced genes were intersected with high-confidence NDD genes from the GeneTrek database v33^34^ to calculate the observed overlap. To obtain a null distribution of the expected overlap, 1317 genes (equal to the total number of alternatively spliced genes upon *Rbfox2*-OE) were randomly sampled from the universe of genes with annotated splice events and the overlap with high-confidence NDD genes and the p-value computed (see Monte Carlo Simulation section for details).

### Monte Carlo Simulation

To evaluate the statistical significance of overlaps for various splicing events and genes, Monte Carlo simulations were performed. In each simulation, a number of genes equal to the observed count of alternatively splicing events (t*Rbfox*-KO and *Ptbp2*-KO) or genes (NDD risk genes) were randomly sampled from the universe of genes or splicing events. The overlap with the gene set of interest (e.g., high-confidence NDD genes) was then calculated. This process was repeated 100,000 times using a seed of 42 to generate a null distribution of expected overlaps. The p-value was calculated as

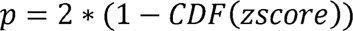

using Python (version 3.10.12)^102^, numpy (version 1.23.5)^100^, and scipy (version 1.10.1)^106^. CDF corresponds to the cumulative distribution function of the normal distribution.

## Data availability

The RNA Sequencing data generated for this study are available on NCBI SRA (Sequence Read Archive) under the accession number PRJNA1013323.

## Acknowledgments

S.W acknowledges financial support from the Emergent Algorithmic Intelligence Center of the Johannes Gutenberg University Mainz funded by the Carl-Zeiss-Stiftung. H.T. and S.G. acknowledge financial support from the Landesinitiative Rheinland-Pfalz and the Resilience, Adaptation, and Longevity (ReALity) initiative of the Johannes Gutenberg University of Mainz. J.W. acknowledges funding from the DFG (WI 3837/10-1).

## Author information

### Contributions

S.W. performed the bioinformatics analyses, interpreted and visualized the results. H.T. contributed to conceptualizing bioinformatics analyses, interpreted the results and performed statistical analyses. A.S. performed cell culture, RT-qPCR, Western blot and Nanopore sequencing experiments. L.S. performed in-utero electroporation experiments and analyzed the data. L.Z. and S.M. performed immunostainings and analyzed the data. S.L.Z. performed Luciferase Assays. A.W. performed preprocessing of Nanopore-seq data. T.V. provided the *in vitro* neuronal differentiation model. DS and SS analyzed data and revised the manuscript. M.H. and S.G. provided supervision and organized funding. J.W. designed the research, supervised the study, and organized funding. S.W., H.T., and J.W. wrote the manuscript. All authors read and approved the final version of the manuscript.

## Ethics declarations

### Ethics approval

All animal experiments were performed in accordance with the regulations of the local authorities (Landesuntersuchungsamt Rheinland-Pfalz, license number 23 177-07/G 13-1-089).

