## Supplementary Figures for "miRNA-mediated expression of RBFOX2 governs the splicing transition from progenitors to neurons in the developing brain"

**Supplemental Figures**

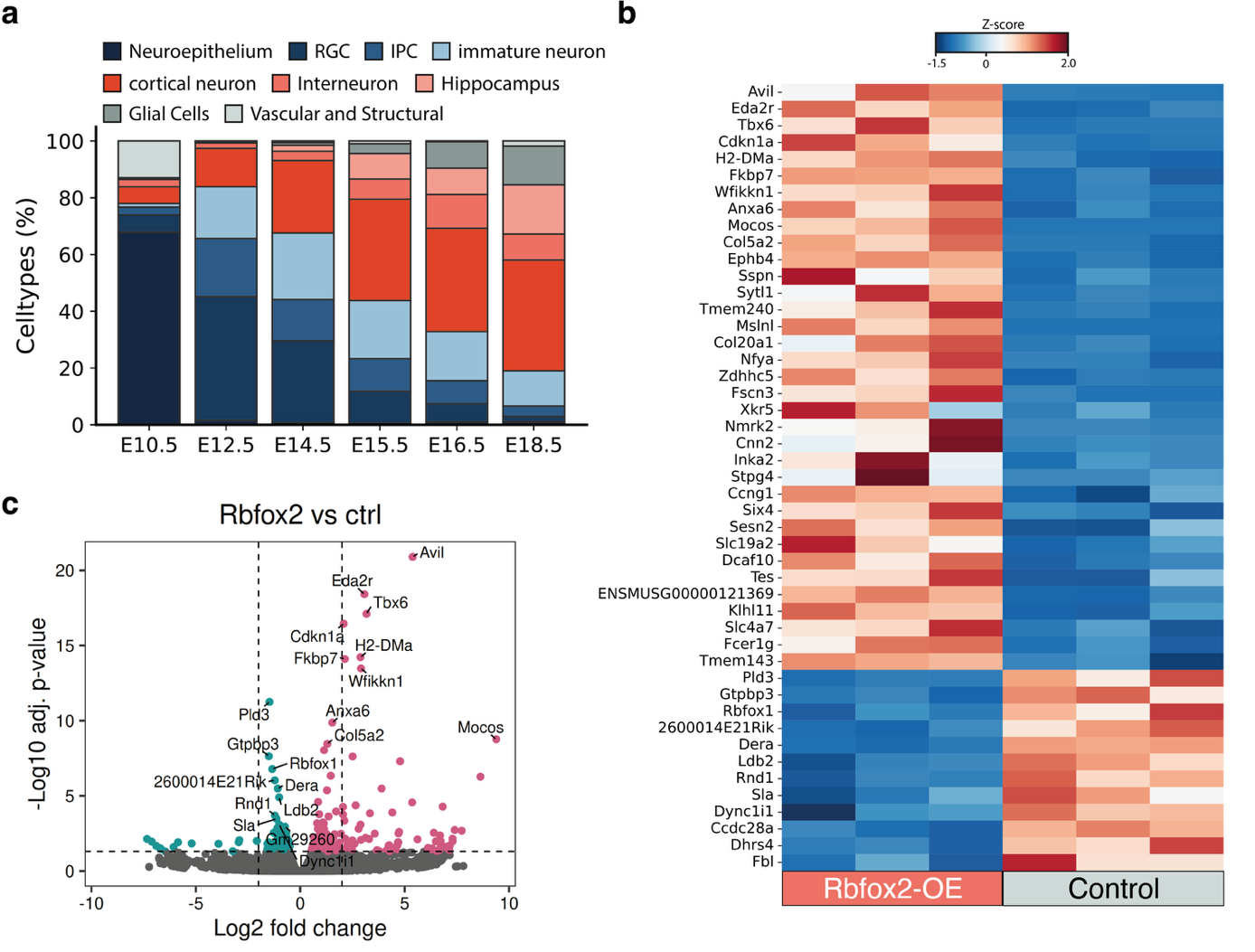

**Supplementary Figure S1 a**) Stacked bar chart showing the relative proportion of different cell types across embryonic ages from E10.5 to E18.5. Cell types are grouped by developmental stage and lineage/differentiation stage: neural progenitors (Neuroepithelium, RGC, IPC) in dark blue shades, immature neurons in light blue, mature neurons (Cortical neurons, Interneuron, Hippocampus) in red/orange shades, and non-neuronal cells (glial Cells, vascular and structural) in gray shades. Early embryonic stages (E10.5-E12.5) are dominated by neural progenitors, while later stages show increasing proportions of mature neurons and glia cells, reflecting the progression of neurogenesis and cortical maturation. Data was obtained from Ruan et al. **b)** Top 50 differentially expressed genes upon Rbfox2-overexpression (Rbfox2-OE) in the embryonic neocortex compared to control samples. Expression values are shown as z-scores. **c)** Volcano plot showing minimal alteration in differential gene expression between Rbfox2-OE and controls.

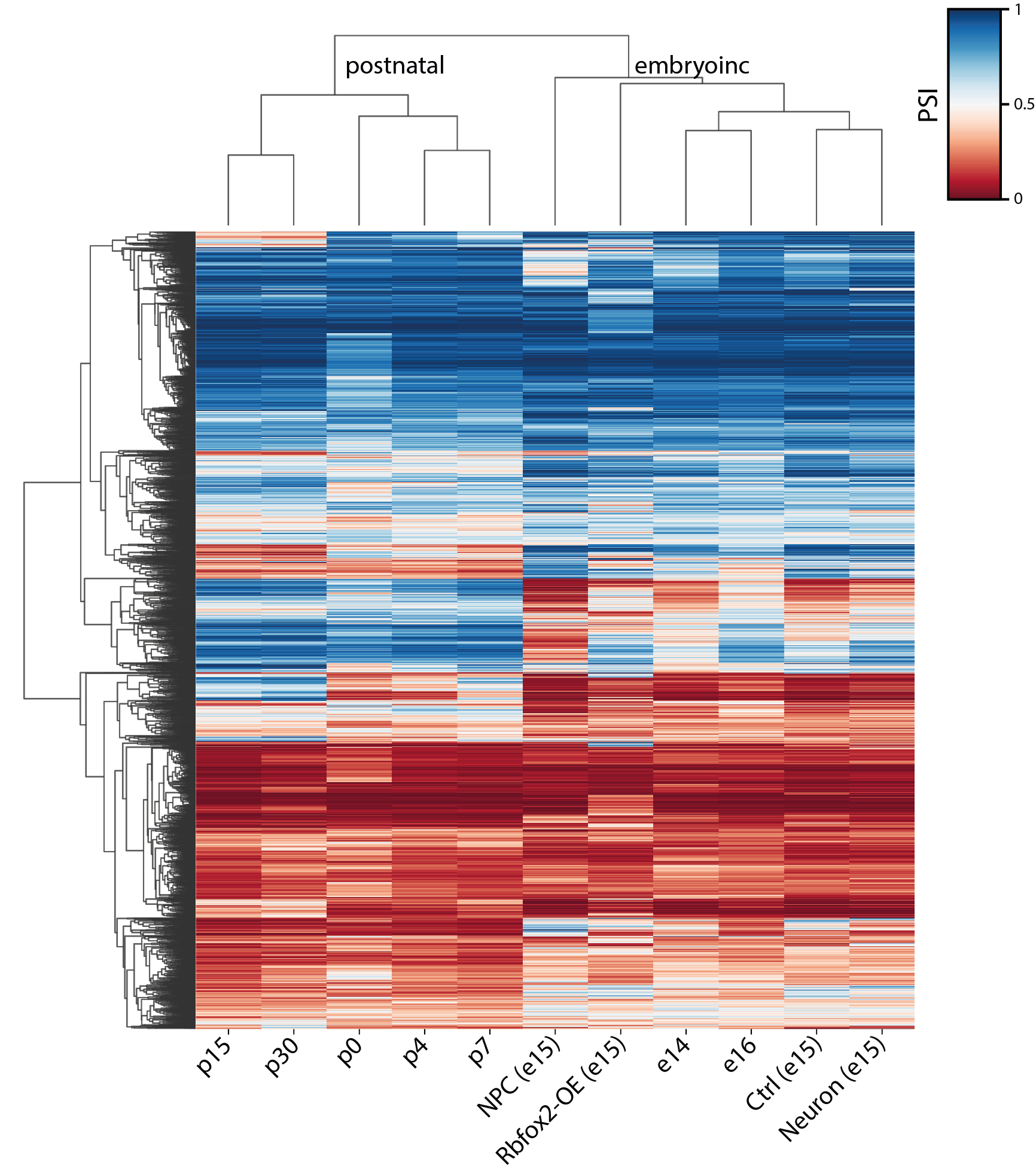

**Supplementary Figure S2** Developmentally regulated splicing events^1,2^ hierarchically clustered using the percentage spliced-in (PSI) values. Embryonic and neuronal maturation stages cluster together. Clearly, Rbfox2 overexpression(OE) leads to an aberrant splicing pattern that does not cluster with samples from comparable developmental stages.

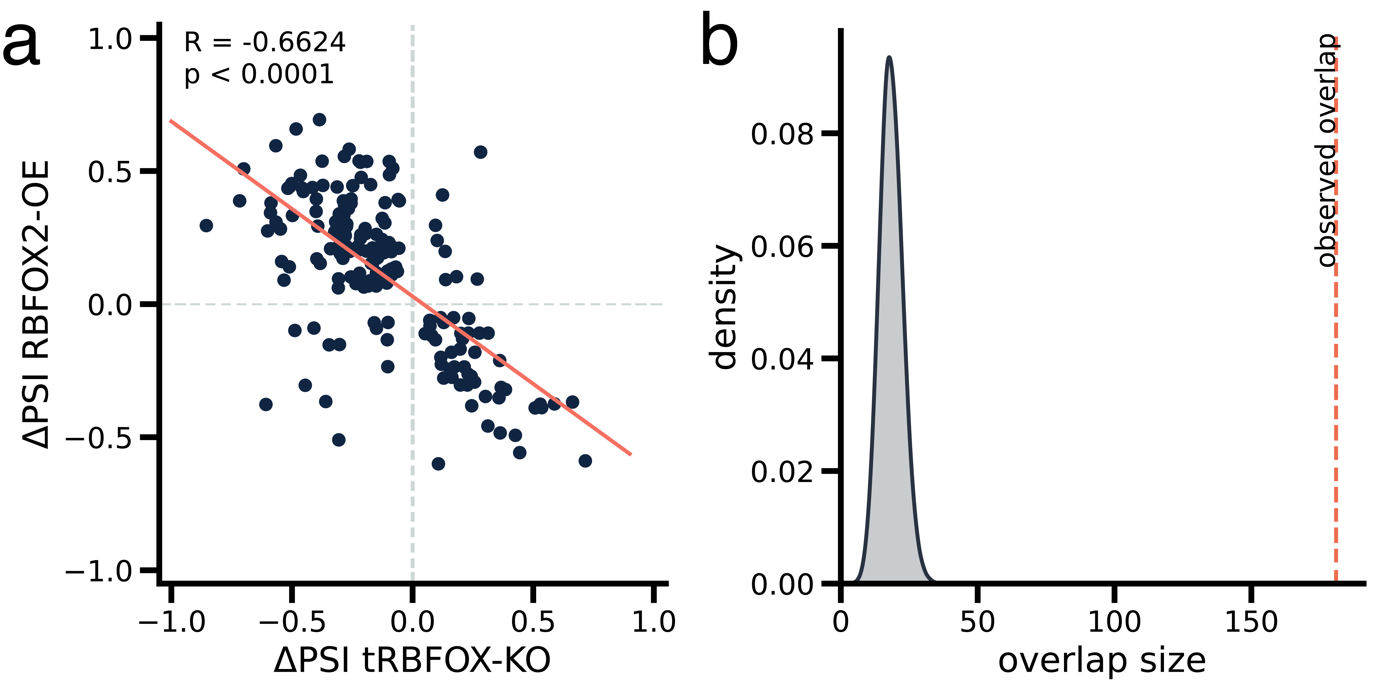

**Supplementary Figure S3** Comparison of splicing changes in Rbfox2 overexpression (OE) and Rbfox triple knockout (tKO) conditions in mouse embryonic stem cells (data from Jacko et al.^3^). **a**) Correlation analysis of splicing changes (ΔPSI) between Rbfox2-OE and tRbfox-KO conditions. Each point represents a significant splicing event detected in both data sets. The x-axis shows ΔPSI values for tRBFOX-KO, while the y-axis shows ΔPSI values for Rbfox2-OE. A strong negative correlation (R = -0.6624, p < 0.0001) confirms the expected opposite splicing effects in OE versus KO conditions. **b**) Monte Carlo simulation results from 100,000 runs, showing the distribution of overlap sizes between Rbfox2-OE and tRBFOX-KO splicing events. The x-axis represents the overlap size, and the y-axis shows the density of occurrences. The observed overlap (indicated by the orange dashed line) is significantly larger than expected by chance (fold change ≈ 9.833, p < 0.0001).

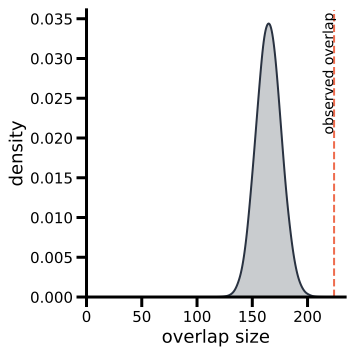

**Supplementary Figure S4** Monte Carlo simulation results from 100,000 runs, showing the overlap distribution between randomly selected genes and high-confidence neurodevelopmental disorder (NDD) genes^4^. The observed overlap between significantly alternatively spliced genes in Rbfox2-overexpression samples compared to controls and NDD genes is significantly enriched (224 out of 1317 genes, fold change ≈ 1.358, p < 2e-07).

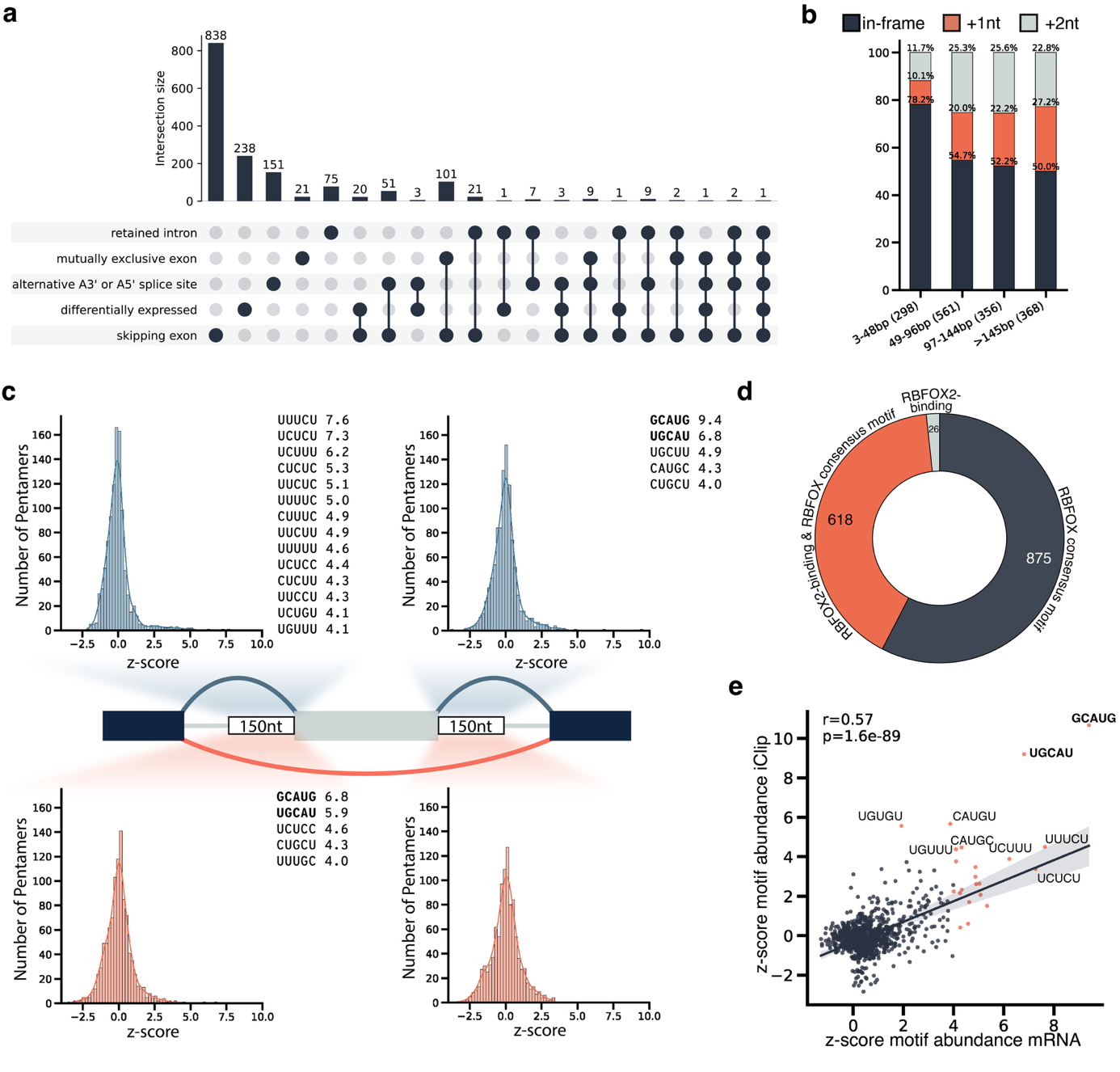

**Supplementary Figure S5** **a)** UpSet plot depicts the overlap between 1) Differentially expressed genes, 2) Genes with significant alternatively spliced skipping exon events, 3) Genes with significant alternatively spliced mutually exclusive exon events, and 4) Genes with other splice events, including retained intron, A3’, and A5’ splice sites between Rbfox2 overexpression (OE) and control samples. **b)** Frequency of induced frameshifts due to alternative splicing for different exon lengths. c) Histogram with the frequency of enriched pentameric motifs around alternatively spliced exons in Rbfox2 OE samples versus control. Blue color denotes motifs associated with exon inclusion, while orange represents motifs linked to exon skipping. Known RBFOX2 motifs are indicated with bold font d) Analysis of alternatively spliced cassette exons reveals that 96% either host an RBFOX motif in the flanking intronic sequence or are bound by RBFOX, as confirmed by iClip data^5^. e) Correlation for position-dependent prevalence of motifs enriched upstream or downstream of alternatively spliced exons in Rbfox2-OE vs. control comparison and in iClip data^5^. Motifs with high correlation are likely recognized by the same splicing factor.

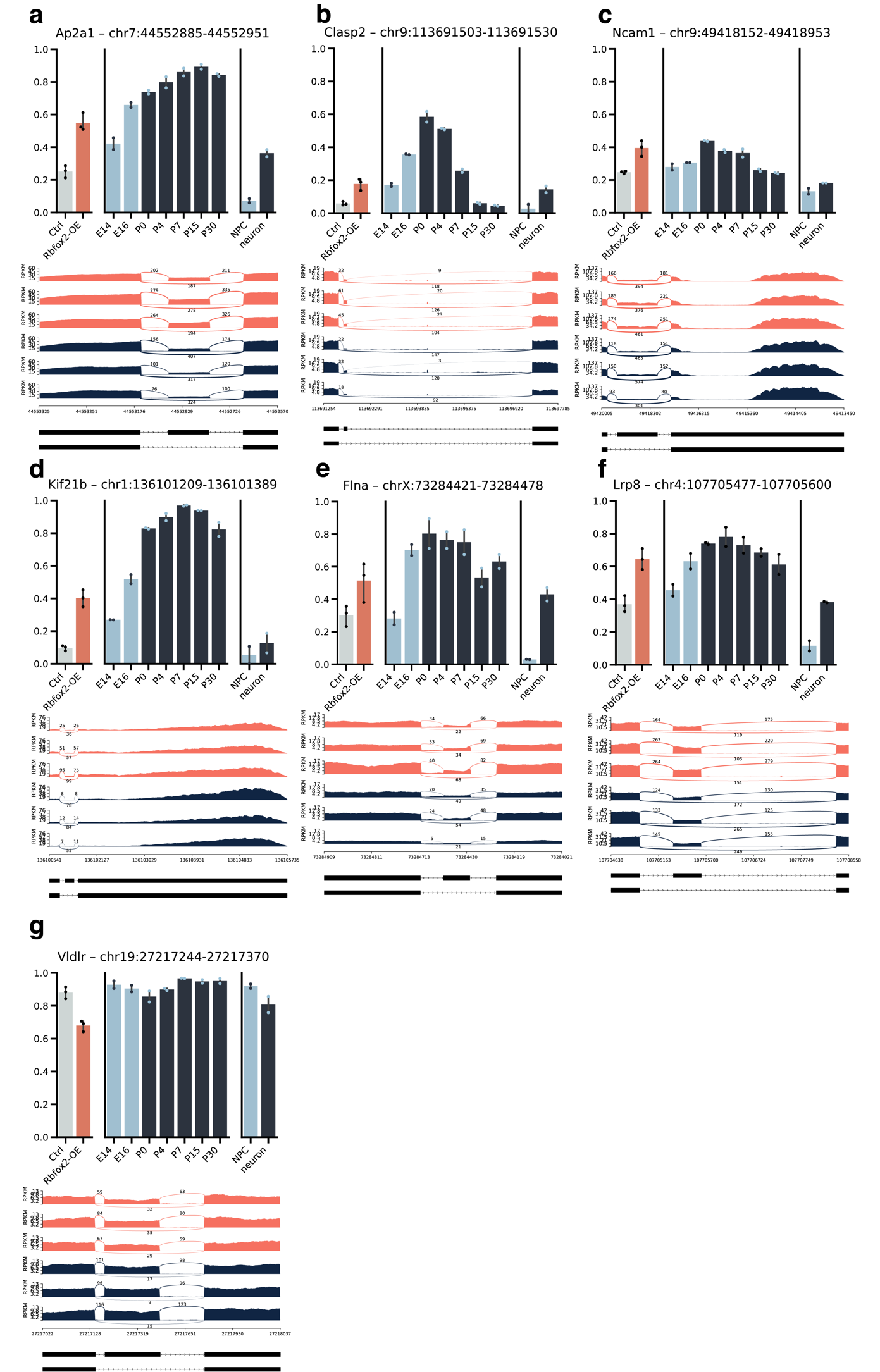

**Supplementary Figure S6** Alternative splicing analysis of representative genes during neocortical development and upon Rbfox2 overexpression. **a-g)** Each panel shows a different gene (Ap2a1, Clasp2, Ncam1, Kif21b, Flna, Lrp8, Vldlr) with bar graphs displaying percentage spliced-in (PSI) values across developmental stages and conditions (top), sashimi plots illustrating read coverage and junction reads (middle), and alternative isoforms from the GFF annotation file (bottom). Bar graphs include control, Rbfox2-OE, embryonic and postnatal stages, NPCs, and neurons. Sashimi plots show three replicates for two conditions (control: blue, Rbfox2-OE: orange).

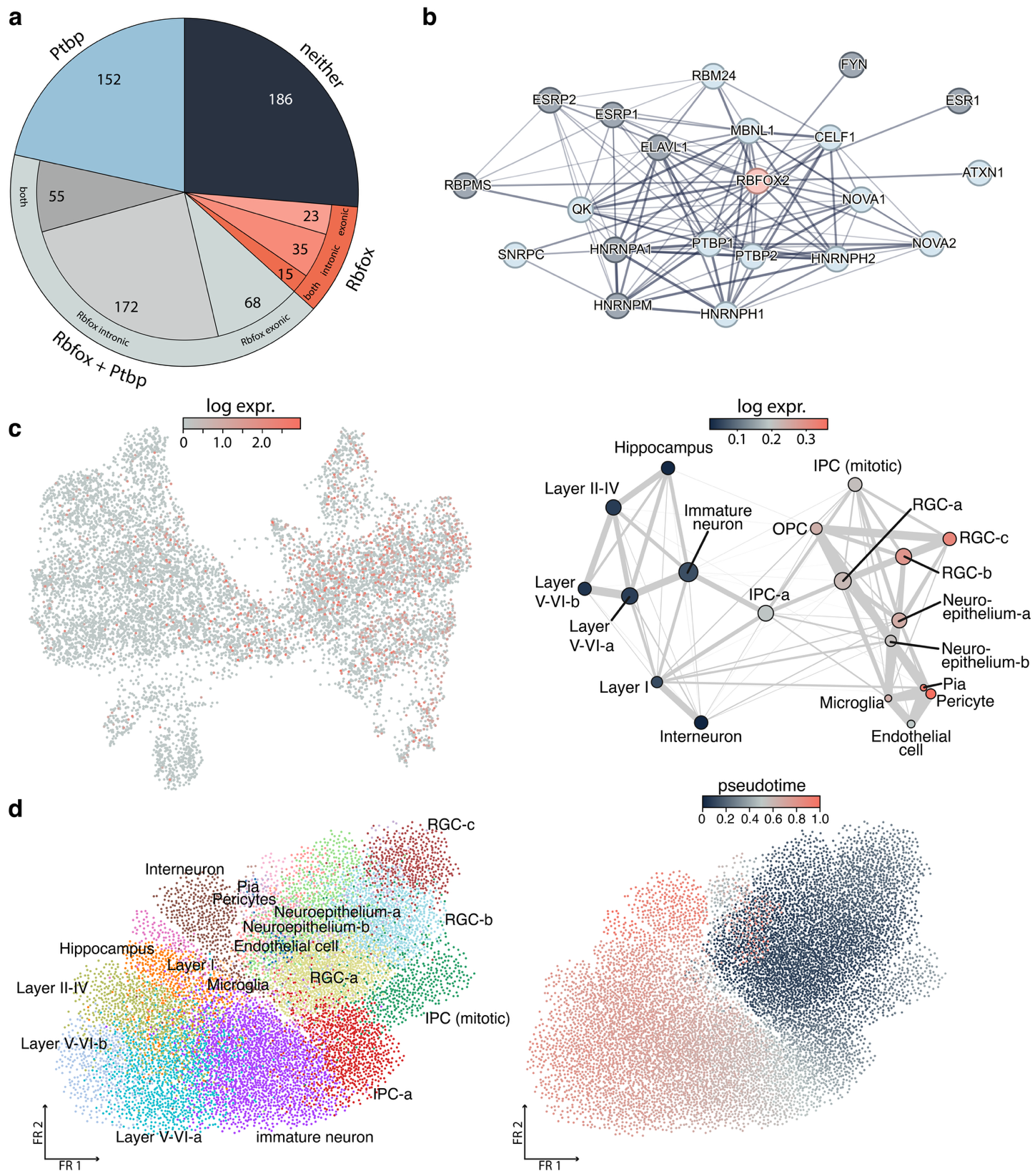

**Supplementary Figure S7** **a)** Analysis of significantly alternatively spliced cassette exons for conserved PTBP and RBFOX motifs. **b)** StringDB protein-protein interaction network showing RBFOX2 (orange) and its interaction partners. High-confidence partners are highlighted in light blue. **c)** Expression of Ptbp1 during neocortical development in different cell types of the embryonic neocortex. Data was obtained from Ruan et al. **d)** Pseudotime computation using the PAGA algorithm (left cluster assignment, right pseudotime).

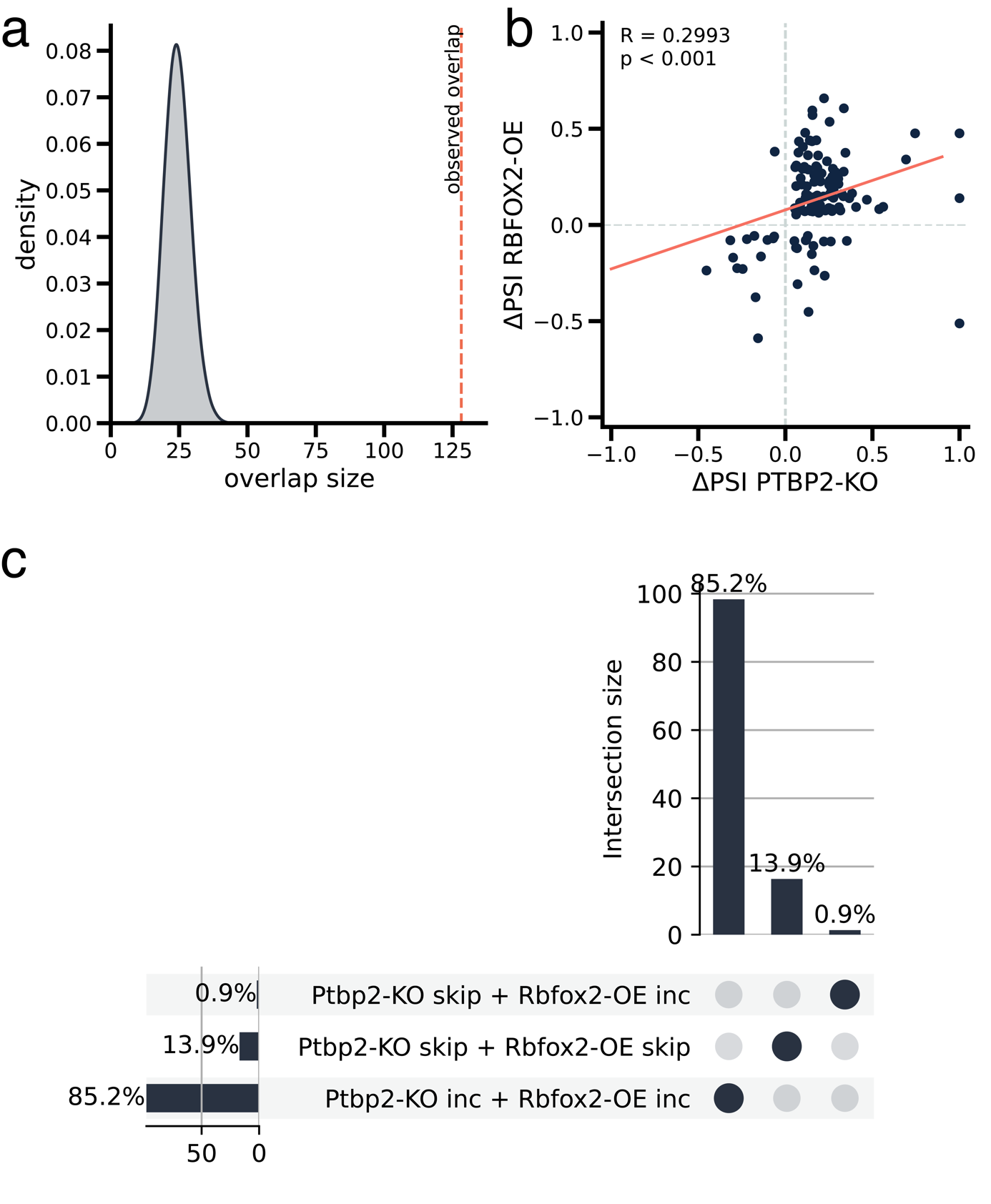

**Supplementary Figure S8** Comparison of splicing changes in Rbfox2-OE and Ptbp2-KO^6^ conditions. **a)** Monte Carlo simulation results from 100,000 runs, showing the distribution of overlap sizes between randomly selected events and those affected by Rbfox2-OE and Ptbp2-KO. The x-axis represents the overlap size, and the y-axis shows the density of occurrences. The observed overlap (indicated by the red dashed line) is significantly larger than expected by chance, suggesting a substantial and non-random relationship between Rbfox2-OE and Ptbp2-KO splicing effects. **b)** Correlation analysis of splicing changes (ΔPSI) between Rbfox2-OE and Ptbp2-KO. Each point represents a splicing event. The x-axis shows ΔPSI values for Ptbp2-KO, while the y-axis shows ΔPSI values for Rbfox2-OE. A moderate, statistically significant positive correlation (R = 0.2993, p < 0.001) is observed, indicating a degree of similarity in splicing effects between the two conditions. **c)** Intersection analysis of splicing events affected by Ptbp2-KO and Rbfox2-OE. The UpSet plot shows the percentages of cassette exons that are included (inc) or skipped (skip) in both conditions. The majority (85.2%) of events show increased inclusion in both Ptbp2-KO and Rbfox2-OE.

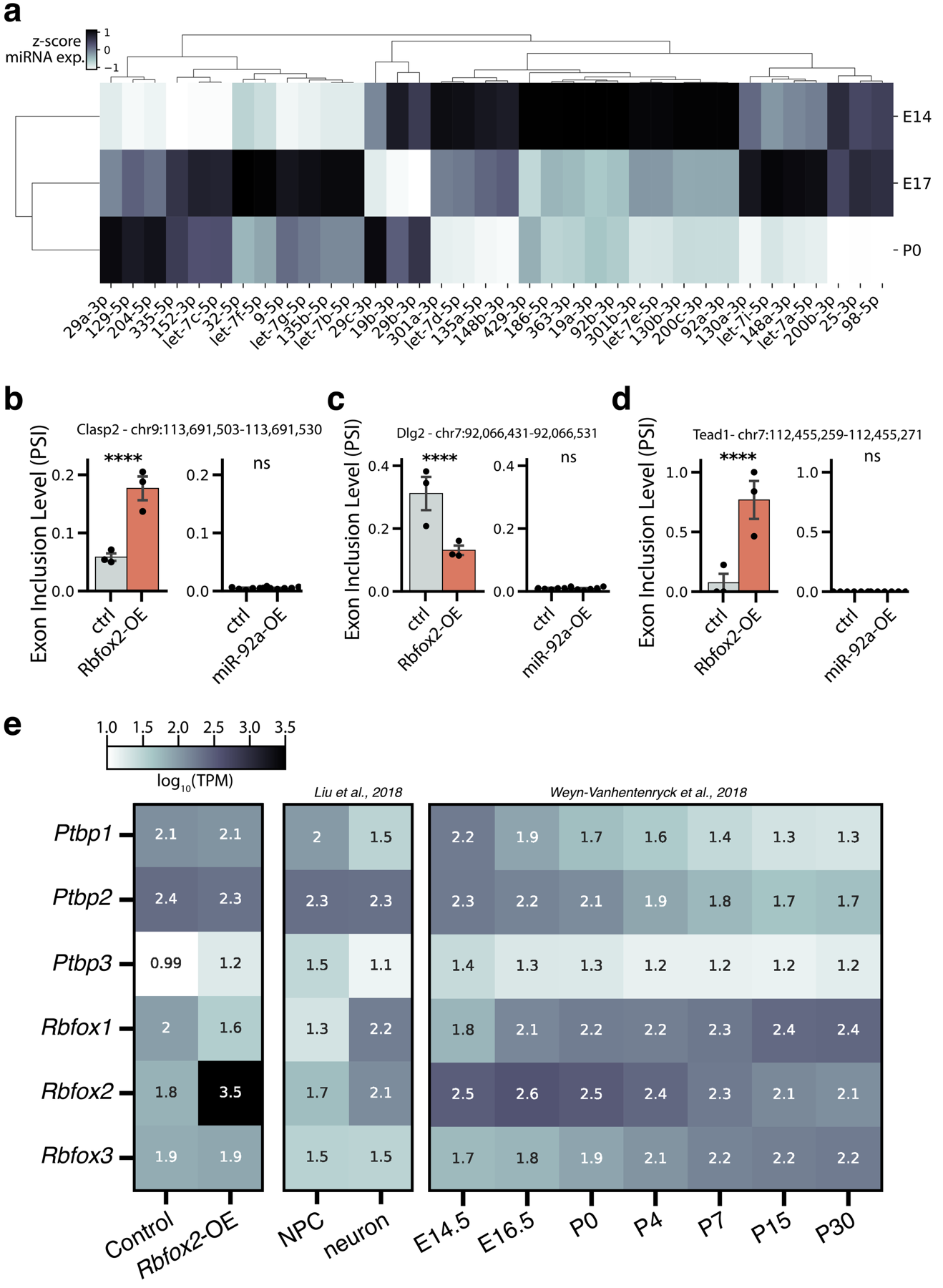

**Supplementary Figure S9** **a)** Heatmap showing the expression of miRNAs predicted to target Rbfox2. Normalized expression values are shown as z-scores. Data was obtained from Todorov et al.^7^. **b-d)** Nanopore long read sequencing of Rbfox2-regulated targets Clasp2 (b), Dlg2 (c), and Tead1 (d) after miR-92a-3p overexpression (OE). Left panel: effect of Rbfox2-OE, right panel: effect of miR-92a-OE. **e)** Heatmap showing the gene expression of the RBFOX and PTBP protein families in our study, as well as in NPCs and neurons from Liu et al. ^8^ and E14.5 to P30 cortical samples obtained from Weyn-Vanhentenryck et al.^1^.Values correspond to log10-transformed transcripts per million (TPM).
